# Associations of waterbirds with harvesting and ploughing events in rice fields: increasing foraging opportunities or just gathering for the feast?

**DOI:** 10.64898/2026.06.29.735228

**Authors:** João Paulino, José Pedro Granadeiro, Edna Correia, Teresa Catry

## Abstract

Surges in food availability create localized but intense foraging opportunities, often attracting multi-species consumer groups. Agricultural practices can trigger these surges, prompting bird associations. However, the strength and duration of the association, as well as its drivers, remain unclear. This study examines waterbird association with harvesting and ploughing events in rice fields. The duration and magnitude of these associations were determined and three hypotheses addressed to explain them: (1) increased food availability, (2) enhanced foraging success and (3) reduced time allocated to vigilance. Waterbird counts and GPS tracking revealed strong associations with management events. Bird numbers spiked during events but declined within one to two days. Food availability (and soil penetrability) increased significantly during events – crayfish and rice during harvesting, worms and soil penetrability during ploughing – supporting Hypothesis 1. However, this did not improve foraging performance (intake rate, foraging success), rejecting Hypothesis 2. Higher competition, interference or kleptoparasitism in these large mixed-species flocks may offset increased food availability benefits. Alternatively, functional responses of target species may limit prey intake due to physiological or behavioural constraints. Hypothesis 3 was also unsupported, as birds did not reduce vigilance. It is plausible that birds may be drawn to events by the perception of a feast, not actual benefits. Gregariousness and foraging behaviour by “local enhancement” may explain such associations. Results highlight the complexity of bird responses to food surges while suggesting waterbirds in rice fields maintain stable foraging performance during agricultural management events and otherwise, indicating resilience to agricultural timing shifts.

## 1. Introduction

Temporary surges in food availability – such as seasonal fruiting (Curran and Leighton 2000), insect emergences (Yang 2004), or nutrient influxes in aquatic systems (Gratton and Denno 2003) – can create brief yet intense foraging opportunities for various organisms (Yang et al. 2008). Beyond these natural cycles, ephemeral but appealing feeding opportunities may also arise from animal or human activities (Catry et al. 2014; Correia et al. 2019; Ouled-Cheikh et al. 2022). For example, predatory fish and marine mammals, during their hunting, drive schools of small fish to the surface, offering concentrated, short-term feeding chances for seabirds like terns and shearwaters (Correia et al. 2019). Similarly, human activities, such as fisheries (Ouled-Cheikh et al. 2022) and agriculture (Catry et al. 2014), generate pulses of resource availability. Fisheries contribute by discarding bycatch or processing waste (Ouled-Cheikh et al. 2022), while agricultural practices like clearing or land preparation expose invertebrates, seeds or crop residues (Lourenço and Piersma 2008; Catry et al. 2014). Regardless of their origin, these superabundant yet temporally and spatially restricted resources often lead to large aggregations of consumers, typically forming multi-species groups of mobile and opportunistic predators (Quérouil et al. 2008; Goyert et al. 2014).

When large numbers of birds are drawn to a single area by the transient resources made available during food surges, behavioural processes play a major role in how these animals derive benefits from the environment (Sridhar et al. 2009; Ye et al. 2017; Zhu et al. 2020). Many of these processes arise from flocking behaviour, as large flocks – often composed of multiple species competing for the same resources – are frequently formed (Sridhar et al. 2009; Ye et al. 2017; Zhu et al. 2020). Mixed-species foraging flocks (Moynihan 1963) often translate into improved foraging efficiency and reduced predation risk (Morse 1977). However, not all participants in flocks necessarily benefit; some individuals may even incur costs from flock association (Zamora et al. 1992; Cimprich and Grubb 1994; Pomara et al. 2007; Catry et al. 2009). While foraging, individuals in groups can use information from others on foraging techniques, foraging locations, or even steal food from other flock members (Stinson 1980; Gonzalez 1996; Yates et al. 2000). Inter- and intraspecific competition within the flock, either direct (kleptoparasitism) or indirect (resource depletion, occupation of foraging space; Ye et al., 2017), can affect food intake rates of individuals (Stinson 1980; Gonzalez 1996; Yates et al. 2000). The intricate interplay between food availability and behavioural changes creates a complex relationship between food surges and bird populations.

The association between birds and agroecosystems – ecological systems shaped by agricultural management – is a well-documented phenomenon (Elphick et al. 2010; Johnson et al. 2011; Catry et al. 2014; Golawski and Kasprzykowski 2021), as these systems often provide abundant and predictable foraging opportunities (Toral and Figuerola 2010; Navedo et al. 2013). However, the level of attractiveness of these areas can be affected by management events (Toral and Figuerola 2010; Navedo et al. 2013). These events consist of discrete, localized short-term interventions in agricultural land that cause immediate and often intense modifications to the environment, many times affecting prey availability and foraging efficiency of predators (Elphick et al., 2010; Ibáñez et al., 2010; Johnson et al., 2011). Changes in vegetation cover or soil structure, for instance, can directly influence the availability, distribution, and structure of food sources (Elphick et al. 2010; Catry et al. 2014), but the impact of these changes is mostly short-lived. For instance, crop harvest in meadows can lead to displacement of insects (Catry et al. 2014) or mammals (Golawski and Kasprzykowski 2021) making them more available to avian predators. Yet, after some time, most prey relocate to more favourable habitat for shelter, sharply decreasing their availability in fields that have undergone the management event (Catry et al. 2014; Golawski and Kasprzykowski 2021). To deal with these ephemeral surges in food, birds typically search for recently managed fields to forage with maximum efficiency (Catry et al. 2014). Although it is widely recognized that birds are often associated with agricultural management events, few studies have directly examined their impact, particularly how long the target area remains attractive to birds and the factors driving this attractiveness.

Rice (*Oryza* sp.) fields are the second largest wetland type worldwide and the largest irrigated crop (International Rice Research Institute 1995; Ramsar Convention Secretariat 2010), with their area increasing in many regions (Yoon 2009; Ramsar Convention Secretariat 2010). Rice fields provide important foraging opportunities for waterbirds, attracting many species especially during the non-breeding season or for short stopover periods during migration (Fasola and Ruiz 1996; Elphick 2000; Lourenço and Piersma 2009; Elphick et al. 2010; Parejo et al. 2019). In the context of worldwide loss of natural wetlands (Ballut-Dajud et al. 2022; Fluet-Chouinard et al. 2023), rice fields are becoming increasingly important for waterbird conservation (Ballut-Dajud et al. 2022; Fluet-Chouinard et al. 2023). Despite providing predictable and abundant food resources, rice fields are very seasonal habitats, undergoing dramatic changes across the annual cycle of rice production and according to different management practices and events (e.g. dry vs. flooded fields, presence/absence of rice plants, etc; Toral & Figuerola, 2010; Navedo et al., 2013). In Europe, rice cultivation is broadly divided into two seasons: the growing season in spring and summer, and the non-growing season in autumn and winter (Blengini and Busto 2009; Nelson et al. 2014). The non-growing season begins in early autumn with harvesting, followed by land preparation – namely ploughing – to create a seedbed for the next year’s sowing (Blengini and Busto 2009). Both these management events attract a large number of waterbirds (Madaleno 2019; Paulino et al. 2025). Moreover, these events are not conducted simultaneously in all rice fields, thus creating a high spatial heterogeneity in environmental conditions within the landscape. This combination of factors provides an ideal scenario to understand how waterbirds respond to management events, in particular to investigate their short-term decisions regarding habitat use and diet and their activity budgets. Such information is essential for evaluating the suitability of rice fields as alternative habitats for waterbirds, especially given the rising concerns about rice crops functioning as ecological traps attracting birds to these dynamic, short-lived habitats, which can become unsuitable for them during significant parts of the year (Elphick 2015).

The objective of this study is to examine the impact of management events in rice fields, specifically harvesting and ploughing events, on the local abundance and foraging dynamics of waterbirds and unveil the drivers of such impact. We have analysed fine-scale temporal patterns in the association of birds with rice fields during management events (using regular counts and GPS tracking), while concurrently measuring changes in bird foraging behaviour and performance, flocking parameters, bird diet and food availability before and after these events. Waterbirds often associate with harvesters and tractors during active harvest and ploughing events in rice fields (Fasola and Ruiz 1996; Ginantra et al. 2023; pers. obs.). This study aims to test three hypotheses for these foraging associations of waterbirds with management events in rice fields: 1) management events increase prey availability; 2) waterbirds associated with these events enhance foraging performance and 3) waterbirds reduce time allocated to vigilance during these events, which can translate into improved foraging budgets.

## 2. Methods

### 2.1. Study area and species

The study area consists of 4700 ha of rice fields located in an agricultural landscape surrounded by large areas of saltmarsh and intertidal mudflats situated in the Tagus river basin (38Ö 57’ N, 8Ö 54’ W) known as Lezíria Grande, near the city of Vila Franca de Xira, Portugal (Fig 1). The southernmost part of Lezíria Grande, including some rice fields, pastures as well as estuarine habitats have been classified as a Natural Reserve and a RAMSAR site since 1976, while a larger area, including most of the remaining rice fields in the study area, have also been classified as a Special Protection Area since 1994 under the network Natura 2000 (Fig 1), due to its biological significance.

**Fig 1.**
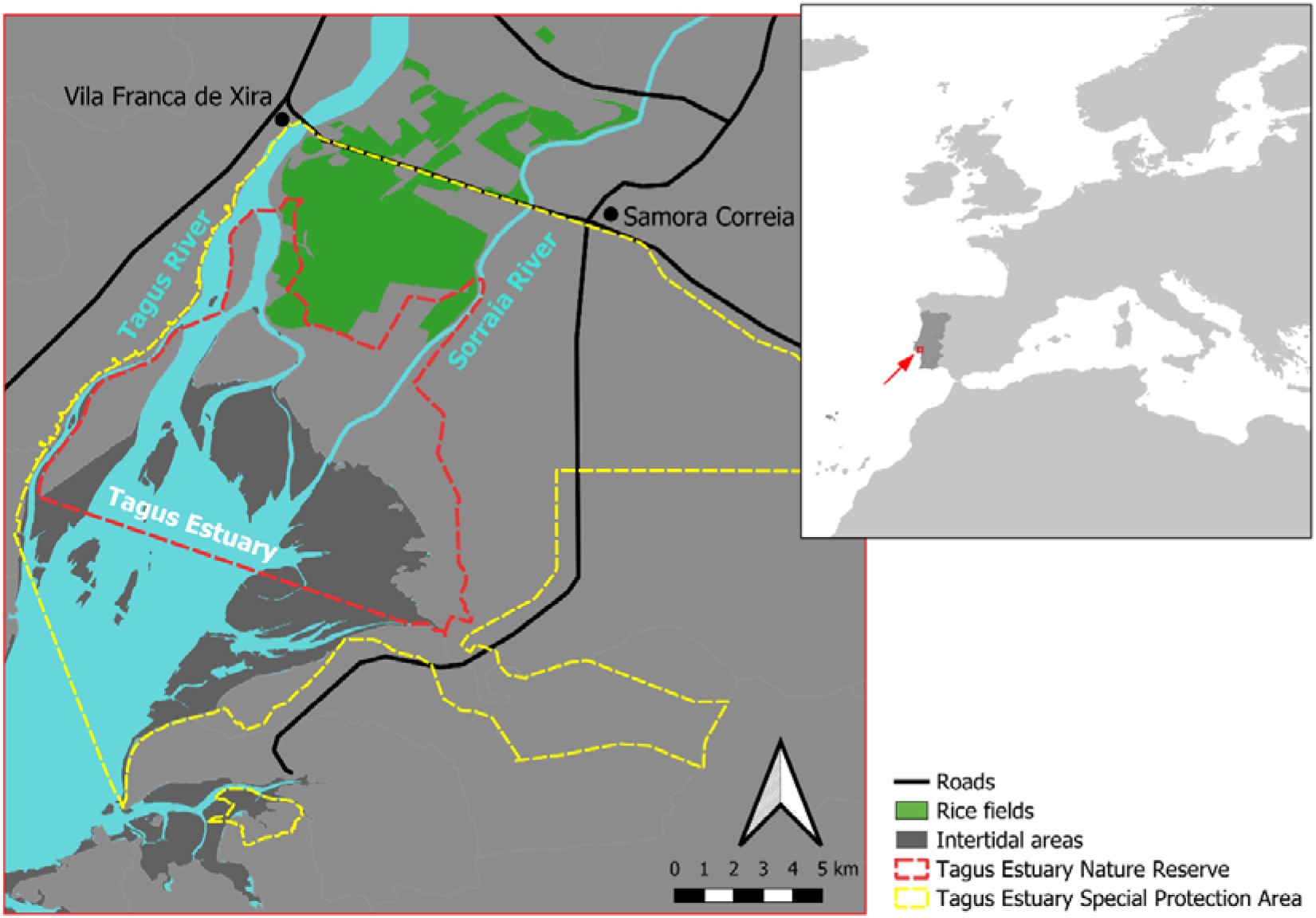
Study area. Map showing the study area including all rice fields in Lezíria Grande.

In the study area, rice is harvested from late September to late November, using a mechanical harvester. Until the new sowing season which occurs from late April to early May, water levels in the rice paddies depend mostly on precipitation (Lourenço and Piersma 2009), although some are intentionally flooded by producers. All paddies are ploughed between November and April. Harvested fields are typically ploughed twice: the first ploughing occurs up to a few weeks after harvest, and the second is closer to the next years’ sowing when the fields are mostly dry. Ploughing in flooded fields is performed with a tractor equipped with paddy wheels and usually a cage roller attached in a process called puddling. In dry fields, ploughing is typically done with a tractor that has rubber wheels and a disc harrow (disking).

This study was conducted between the end of the rice growing season (August) and the end of the non-growing season (March) in 2021-2022 and 2022-2023.

The present work focused on waterbirds – bird species that are ecologically dependent upon wetlands (Wetlands International 2012) – using rice fields during the non-growing season. Although the waterbird community as a whole was studied, special emphasis was placed on two waterbird species – glossy ibis (*Plegadis falcinellus*) and white stork (*Ciconia ciconia*) – due to their winter-round presence in the study area (Paulino et al. 2025), their known association with management events (Madaleno 2019; Paulino et al. 2025) and their suitability for research studies (large body size, flocking behaviour, and easy detection and observation to collect data on foraging performance). The number of wintering glossy ibis has significantly increased since 1994, especially in the estuaries of the Tagus and Sado rivers, mostly associated with rice fields, with an estimated 8,320 birds recorded in 2015 (Encarnação 2019) and over 20,000 in 2022, making it one of the most abundant waterbirds in rice fields (Paulino et al. 2025). The white stork population has been growing in the Iberian Peninsula in recent decades, particularly the number of wintering individuals (Encarnação 2015). This increase is driven by factors such as greater food availability due to the spread of crayfish, the presence of landfills, and milder winter temperatures (Catry et al. 2017). During winter, white stork is most commonly found in rice fields and landfills (Catry et al. 2017).

### 2.2. Waterbird counts and GPS tracking

To study the magnitude and temporal patterns of birds’ association with management events (harvest and ploughing), we conducted regular waterbird counts and tracked the movements of glossy ibis using GPS-GSM transmitters. Waterbird counts were performed in 51 different paddies that underwent harvest events and 23 that underwent ploughing events. Harvest and ploughing events were always separated by at least one month. Of the 23 ploughing events studied, one occurred in a paddy that had previously been monitored during harvest events. The latest harvest and earliest ploughing evens studied (regardless of location) in a given season were separated by at least one month. When selecting paddies for study, paddies adjacent to those that had been managed within the previous week were avoided. Counts were performed in the day before (day -1) the management event (either harvest or ploughing), during the event (day 0) and in the 1^st^, 2^nd^, 7^th^ and 14^th^ day after the event. Therefore, each paddy was counted six times in the span of 15 days. Waterbird counts were performed from a stationary point using binoculars (10× magnification) and a zoom-in telescope (20-60× magnification) whenever necessary. In this study, we considered waterbirds all birds belonging to the orders Anseriformes, Ciconiformes, Charadriiformes, Galliformes, Gruiformes, Pelecaniformes and Phoenicopteriformes. Some bird species belonging to these orders that are usually not considered waterbirds (e.g. cattle egret, *Bubulcus ibis*, northern lapwing, *Vanellus vanellus*, etc) have also been included in the study due to their strong association with freshwater habitats in the study area. During the counts, the flooding regime (dry or flooded) of the rice paddies was recorded.

Glossy ibis were captured with mist-nets at rice fields between August and November in 2022 and August and October in 2023. Three different models of solar powered GPS-GSM transmitters (Ornitela, UAB) were used in this study: OrniTrack-9 (9 grams), OrniTrack-10 (modification with elevated solar panel, 12 grams) and OrniTrack-15 (modification with elevated solar panel, 17 grams). Trackers were fitted using a leg-loop harness with a 2 mm Teflon ribbon or using a backpack harness with a 6 mm Teflon ribbon. The weight of the transmitter and the harness represented less than 3% of the bird body mass.

Transmitters were programmed to collect a GPS fix (alongside the timestamp and instantaneous speed) every 20 minutes during daytime and every 2-4 hours during the night period. Fix-interval during the day was increased to 2-hours whenever the battery level decreased below 50%. Tracking data were filtered to match the harvesting period at the study area (25 September to 20 November in 2022, 17 September to 16 November in 2023), but this period also includes ploughing in part of the rice fields. Since glossy ibis forages almost exclusively during the day (pers. obs.), an additional filter was applied to exclude GPS locations from the night period, using *RchivalTag* package (Bauer 2023) in R to identify locations between sunset and sunrise. We have also removed all GPS fixes with instantaneous speeds higher than 10 km/h to exclude flight locations.

To measure the proportion of ibis GPS locations associated with harvesting and ploughing events in rice fields, the locations of all harvesters and ploughing tractors as well as the “status” of each rice field (harvest and ploughed vs non-harvested and non-ploughed) were recorded during regular (every 1-5 days) car transects covering the whole study area during the harvesting season. In addition, to get a more accurate daily “status” of all rice fields, we visually inspected aerial images using Sentinel-2 imagery available at the Copernicus Browser (https://dataspace.copernicus.eu/browser/). For each day during the harvesting period, each rice field was classified as “undergoing management event”, “no event” or “unknown”. Fields were classified as “undergoing a management event” if a harvester or tractor was operating in the field on the day of observation or if the observation occurred within two days after the operation. This time interval was chosen based on the results of bird counts, corresponding to the period after the management event in which birds were still recorded foraging in the fields, i.e., the duration of the association. We then crossed this information with the GPS locations of tagged ibises and calculated the mean proportion of overlap with management events across the study period. To account for the variability in the number of fixes among individuals and days, we first estimated daily proportions of overlap with management events for each bird and then averaged these proportions to obtain a population metric. For 2022 we obtained a total of 8145 GPS fixes from 11 individuals (mean number of birds per day = 9.7) and for 2023 we obtained 12288 GPS fixes from 20 individuals (mean number of birds per day = 15.5). These GPS fixes include overlaps with field status classified as “undergoing management event” and “no event”. Points overlapping fields with status classified as “unknown” were excluded.

Manly selection ratios were calculated using the ‘widesII’ function in the package *adehabitatHS* (Calenge and Basille 2024) to assess the relative daily preference of glossy ibis for fields undergoing management compared to its availability. Field availability was measured as the area of rice fields. Manly selectivity ratios with 95% confidence intervals that do not overlap 1 indicate significant preference (>1) or avoidance (<1).

### 2.3. Waterbird diet

To determine the diet of waterbirds in rice fields at different management stages, we collected droppings from glossy ibis and pellets from white stork throughout the study period. These species were chosen since they are some of the most abundant in the area, making up 23% and 5% of the total number of birds using the studied rice fields over the year (Paulino et al. 2025). Moreover, these species are known to associate with management events (Madaleno 2019; Paulino et al. 2025). Samples were collected after observing individual birds producing droppings and pellets before the harvesting season (August-September; when no fields had been harvested), during the harvesting season (October-November; within fields that had been harvested) and during the ploughing season (December-April; in fields that had been ploughed). It is important to note that during the harvest season, some fields were already being ploughed. Likewise, when ploughing begins, there were many harvested fields. Since the exact location where the birds fed before producing droppings or pellets is unknown, it is possible that some food items in the samples may have been ingested in fields with different management practices. All samples were stored frozen until laboratory analysis. In the laboratory, the samples were examined under a binocular magnifier (10-150× zoom) and food remains were identified to the lowest possible taxonomic level, using a reference collection of invertebrates and rice collected at the study area.

To assess the abundance of two key food items in the diet of waterbirds, all crayfish (*Procambarus clarkii*) and rice grains were counted. The number of crayfish was estimated by either pairing claws, mandibles, or zygocardiac ossicles of similar size, or by counting the number of urocardiac ossicles, whichever method yielded the largest number of individuals. The absence of these structures in samples with the presence of other remains such as carapace debris, precluded the estimate of the number of individuals, so at least one individual was considered in these cases. The number of rice grains in a sample was determined by counting the number of radicles.

The importance of each prey type in the diet of various waterbird species across seasons was evaluated using the frequency of occurrence, defined as the proportion of droppings or pellets containing a given prey taxon. For crayfish and rice, the number of items per dropping/pellet was also determined across seasons.

### 2.4. Food availability

To investigate temporal changes in food availability in paddies that were subjected either to harvesting or ploughing, potential food items were sampled in selected paddies, following the same calendar used for waterbird counts (days -1, 0, 1, 2, 7 and 14).

Crayfish was sampled by walking along 50-meters-long and 5-meters-wide linear transects within the rice paddies (at least five meters from the border of the paddy) and counting all live individuals. Five transects were travelled in each paddy in each time period.

Benthic macroinvertebrates – invertebrates >0.5 mm living on or within the benthic zone, that is, the surface and subsurface layers of sediment at the bottom of a water body – and rice kernels were sampled using sediment cores (28.27 cm^2^, ca. 5 cm deep) collected at random points in each of the selected paddies at least five meters away from the paddy edges and 10 meters away from each other. After performing some test sampling with 20 cm deep cores, it was concluded that most small macroinvertebrates and rice grains (≈98%) where found in the upper layers of the sediment, and therefore only the top 5 cm of sediment was collected and sieved subsequently through a 0.5 mm mesh size, following established protocols (Lourenço et al. 2017; Herbert et al. 2018; Paulino et al. 2021). Five sediment cores were collected in each paddy in each sampling date. All invertebrates and rice kernels were immediately stored in ethanol 96% until further analysis. In the laboratory, all invertebrates were identified to the lowest possible taxonomic level, using guides for the most common families of freshwater invertebrates (Serra et al. 2009). All plant material that could not be confidently identified as rice was classified as “unidentified plant material,” including small seeds (<1 mm) and miscellaneous plant matter. All food items were counted.

Since food availability depends on both the abundance of food items and their accessibility (Nam et al. 2015), we also measured soil penetrability – used as a proxy for accessibility – in fields under different management conditions (rice growing, harvested, and ploughed) and flooding regime (dry or flooded). Penetrability was measured by recording the penetration depth of a 53 cm long, 2 kg iron spike dropped from a height of 1 meter into the sediment. After conducting 180 test measurements in five rice fields, it was determined that penetrability did not change significantly over time within the same management type (Online Resource 1). Therefore, additional samples were collected based solely on field management, without considering the time elapsed since the event. The spike was dropped at random locations within the selected paddies, except in harvested fields. Harvested fields often exhibit deep ruts caused by the tracks of the harvester. The sediment within these ruts has different properties compared to non-rut sections. As approximately half of the harvested field area becomes rutted after harvesting, half of the penetrability measurements were conducted within the ruts, while the other half were taken in non-rut areas.

Waterbird counts performed during this study revealed that management events in dry fields attracted a very low number of birds (see Results). Therefore, since the focus of the study was related with the effects of management events on bird numbers, these fields were not included in further analyses and were not sampled for food availability.

### 2.5. Foraging behaviour and performance, activity budgets and composition of mixed-species flocks

To evaluate the impact of management events on the foraging success and the time allocated to vigilance behaviour of waterbirds, we recorded 2- to 3-minute videos of individual birds foraging in fields at various management stages (before harvest, during harvest, after harvest, during ploughing, and after ploughing). The “after harvest” and “after ploughing” stages were defined as at least three days following the respective management events in a given field. These thresholds were based on the patterns of field attendance by birds as revealed by the waterbird counts – the number of birds decreased dramatically after the second day following the management event (see Results). To ensure the same species were observed across all stages, two of the most abundant waterbird species in rice fields were selected: glossy ibis and white stork.

Video recordings were analysed using the software VLC media player (3.0.20 Vetinari) to collect data on (1) intake rate (number of successful feeding attempts, i.e. attempts resulting in the capture and swallowing of a food item, per minute), (2) foraging success rate (proportion of foraging attempts, i.e. each time a bird penetrated the sediment with its beak, that resulted in successes), (3) foraging effort (number of steps per minute) and (4) time allocated to vigilance behaviour (proportion of time when foraging was interrupted, and the bird raised its head).

To assess how bird behavioural responses influence whether and how waterbirds benefit from management events, the activity budgets and flock parameters of waterbird mixed-species foraging flocks, containing either glossy ibis or white stork, were examined. To determine the activity budgets of both species – specifically, the proportion of time spent foraging versus non-foraging – opportunistic counts were conducted during fieldwork targeting selected paddies with flocks of either species. A total of 214 opportunistic counts were conducted across 46 distinct paddies, encompassing 29 different fields. During these counts, the proportion of individuals engaged in foraging activities in the paddies was recorded. Observations were made during the same management classes used for video recordings: harvest, after harvest, ploughing, and after ploughing. These flocks were further characterized in terms of mixed-flock parameters by recording the total size of the flock, the proportion of conspecifics (ibis or storks) within the flocks, the proportion of heterospecifics, and the total number of the most abundant species present in the flocks (glossy ibis, white stork, lesser black-backed gull, Black-headed gull and Cattle egret) across different management stages.

### 2.6. Data analysis

To investigate differences in the mean number of waterbirds and the mean density of food items among sampling dates (days -1, 0, 2, 7 and 14 in relation to the day of management events) during both the harvesting and ploughing seasons, we performed generalized linear models (GLM) with negative binomial family, followed by Tukey post hoc tests with adjusted p-values for multiple comparisons. GLMs were also used to test differences in parameters collected from video recordings (mean food intake rate, foraging success rate, foraging effort, and time allocated to vigilance behaviour) among periods before and after management events (see above), and in mean soil penetrability under varying rice field management practices and flooding regime (dry or flooded). A beta family was used for parameters expressed as proportions, while all others were tested using a negative binomial family. The same method was applied to test for differences in parameters related to activity budgets and composition of foraging flocks gathered through opportunistic counts across various management stages. These parameters included the proportion of glossy ibis or white stork engaged in foraging activities, the average number of conspecific ibis or storks in targeted mixed-species flocks, the proportion of heterospecifics in mixed-species flocks containing storks or ibises and the total number of different species within the mixed-species flock.

All analyses were performed using R (R Core Team 2025).

## 3. Results

### 3.1. Magnitude and temporal patterns of waterbird associations with harvest and ploughing

A total of 51 harvest events (27 in flooded fields and 24 in dry fields) and 23 ploughing events (all in flooded fields) were counted. Eight waterbird species were counted in the studied rice paddies during the harvesting period. All eight species were observed in flooded fields, but only three were found in dry fields (Online Resource 2). The most abundant species in flooded fields were the lesser black-backed gull, the glossy ibis, the white stork, and the Black-headed gull while the dominant species in dry fields was the Cattle egret (Online Resource 2). From the 15 species recorded during the ploughing period, the more abundant were the glossy ibis, followed by the Black-headed gull and the lesser black-backed gull (Online Resource 2).

We found a significant effect of both harvesting and ploughing events in the abundance of waterbirds, translated into an increase in bird numbers between the counts on day -1 (before management event) and the counts on day 0 (right after the management event) (Fig. 2, Online Resource 3 a-c). The number of birds declined significantly in the following days, and one week after harvest in flooded fields, almost no birds were recorded (Fig. 2a and b). In dry fields, the decline was even more pronounced, with almost all birds disappearing the day after the harvest event. A similar pattern was observed in ploughed paddies, where bird numbers declined significantly just one day after the event, remaining stable throughout the following week and showing a further significant decrease in the second week after ploughing (Fig. 2c). However, not all species responded similarly. Greater flamingos showed a brief increase one day after ploughing before declining, whereas Black-tailed godwits peaked about one week post-ploughing – when few other waterbirds remained – and disappeared by week two (Online Resource 2).

**Fig 2.**
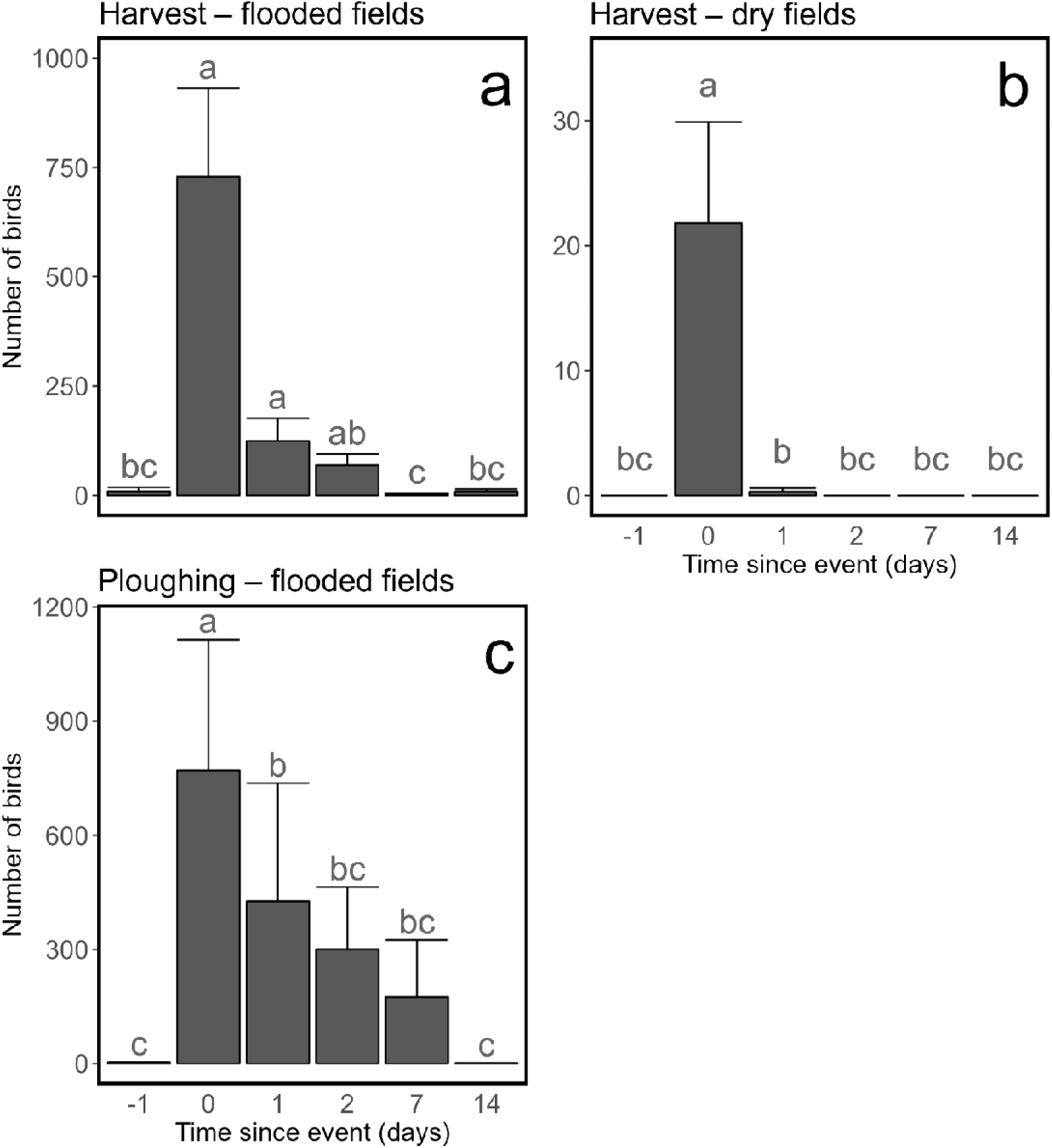
Mean number of waterbirds counted in rice paddies in selected days before, during, and after harvesting and ploughing events. Error bars represent the standard error (SE). a – Total number of birds in flooded harvested fields (n=27); b – Total number of birds in dry harvested fields (n=24); c – Total number of birds in flooded ploughed fields (n=23). On the y-axis, “day 0” refers to the count performed immediately after the management action (harvesting or ploughing) has finished. Differences in the mean number of birds among counting days were assessed using GLMs with negative binomial family followed by Tuckey post hoc tests (with adjusted *P-*values for multiple comparisons) and significant differences (*P*<0.05) are depicted by bars (days) not sharing the same letters.

A total of 27 Glossy ibis where GPS tagged with three different models of solar powered GPS-GSM transmitters (Ornitela, UAB): 5 with OrniTrack-9 (9 grams), 21 with OrniTrack-10 (modification with elevated solar panel, 12 grams) and one with OrniTrack-15 (modification with elevated solar panel, 17 grams). During the harvesting season, 66.1% (± 3.4% SD, n=10 individuals) and 62.4% (± 13.1% SD, n=20 individuals) of the GPS fixes of glossy ibis overlapped with rice fields undergoing a management event, respectively in 2022 and 2023 (Fig 3). This sharply contrasts with an average of only 13.2% (± 6.9 SD) and 11.7% (± 6.8 SD) of the fields having a management event each day. Therefore, Manly selectivity ratios (with 95% CI) showed that glossy ibis have a significant preference for actively managed rice fields in 90% of the days in 2022 and 81% in 2023 (Online Resource 7).

**Fig 3.**
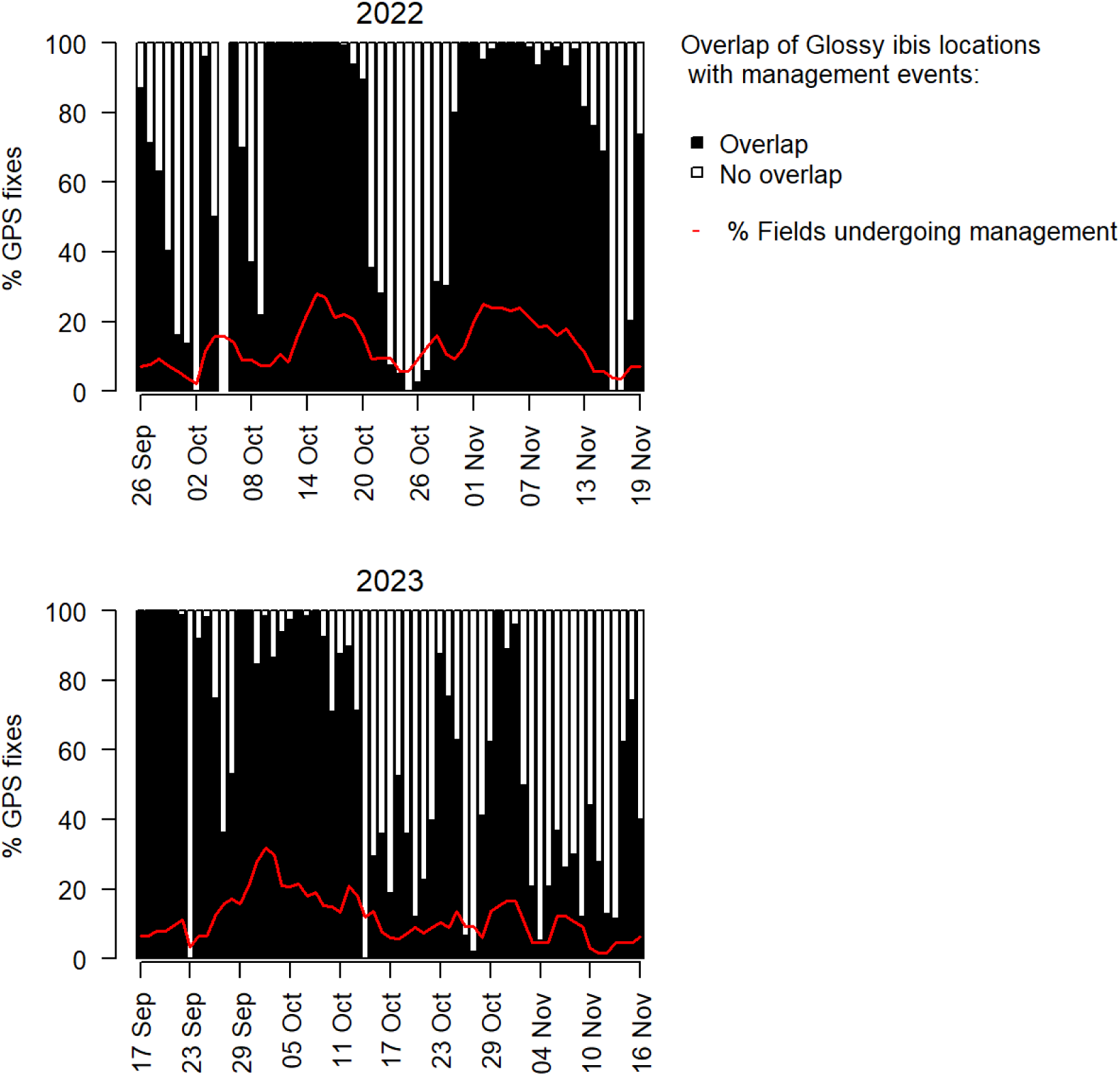
Overlap of glossy ibis GPS fixes with management events. Percentage of GPS fixes of glossy ibis in rice fields undergoing a management event, i.e., during harvesting or ploughing events (overlap), or fields not actively managed (no overlap). Fields were classified as “undergoing management event” whenever there was a harvester or a tractor working in the field in that day or in the two following days. Daily proportions are the mean proportions per bird, to account for the number of different fixes among individuals (mean of 9.7 birds per day in 2022 and 15.5 in 2023; see methods for further details). The red line represents the mean proportion of fields undergoing a management event.

### 3.2. Waterbird diet

Diet composition of the three studied species, as represented by the frequency of occurrence (FO) of identified food items, is summarized in Figure 4. Overall, crayfish and rice were the most frequently consumed food items.

**Fig 4.**
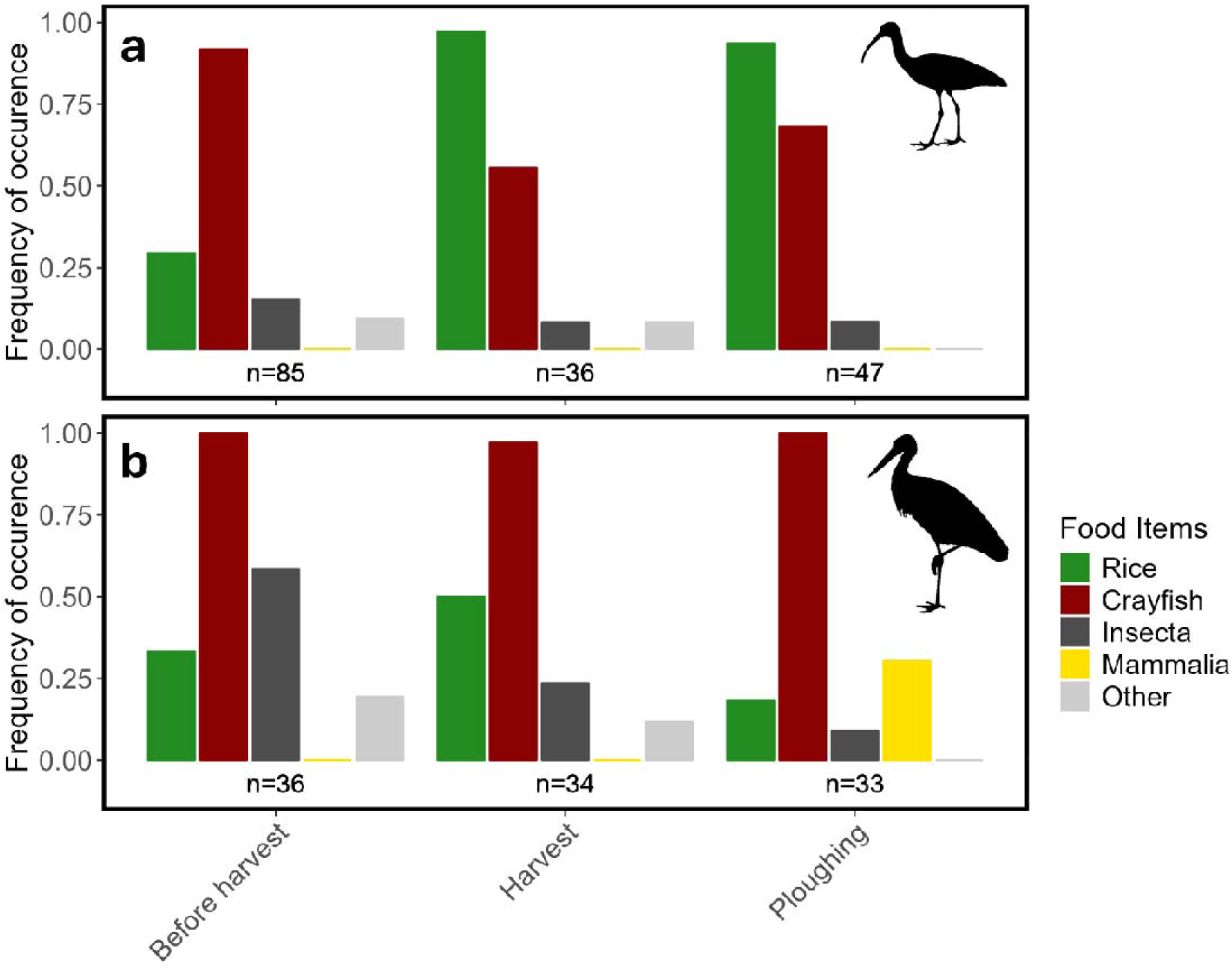
Waterbird diet. Diet composition of (a) glossy ibis (droppings) and (b) white stork (pellets) estimated as the frequency of occurrence of prey items. The food item class “Other” includes unidentified plant material, Arachnida, Bivalvia, Clitellata, Gastropoda and Actinopterygii.

The diet of the glossy ibis (Fig 4a) was dominated by crayfish before harvest (FO = 0.91), with rice as the second most frequent item (FO = 0.29). During the harvest and ploughing seasons, these two items remained the primary components of their diet, but rice became the most frequently consumed (FO harvest = 0.97; FO ploughing = 0.94), followed by crayfish (FO harvest = 0.56; FO ploughing = 0.68). During harvest, the number of rice grains in glossy ibis droppings increased significantly and remained consistently high throughout the ploughing period (Table 1, Online Resource 4a). In contrast, the number of crayfish per dropping remained stable year-round (Table 1, Online Resource 4b).

**Table 1.** Mean (±SD) number of rice grains and crayfish individuals per dropping of glossy ibis and pellet of white stork in three different management stages. Differences in the mean were assessed using GLMs with negative binomial family, followed by Tuckey post hoc tests (with adjusted *P-*values for multiple comparisons) and significant differences (*P*<0.05) are depicted by values in the same line not sharing the same letters in superscript.

| Waterbird species | Food item | Number of items per sample |  |  |
| --- | --- | --- | --- | --- |
|  |  | Before harvest | Harvest | Ploughing |
| Glossy ibis | Rice | 2.11<sup>b</sup><br>(±5.53; n=85) | 9.22<sup>a</sup><br>(±8.47 n=36) | 11.4<sup>a</sup><br>(±16.0; n=47) |
|  | Crayfish | 0.95<sup>a</sup> | 0.56<sup>a</sup> | 0.72<sup>a</sup> |

|  | * | (±0.31; n=85) | (±0.50; n=36) | (±0.54; n=47) |
| --- | --- | --- | --- | --- |
| White stork | Rice | 0.83<sup>a</sup><br>(±1.58; n=36) | 1.35<sup>a</sup><br>(±1.87; n=34) | 0.21<sup>b</sup><br>(±0.48; n=33) |
|  | Crayfish | 18.0<sup>a</sup><br>(±14.7; n=36) | 15.9<sup>a</sup><br>(±12.9; n=34) | 8.03<sup>b</sup><br>(±5.13; n=33) |

\*most crayfish remains were found primarily as carapace debris in glossy ibis droppings, precluding an accurate estimate of the number of individuals and typically leading to an estimate of one individual per dropping.

The diet of white stork was consistently dominated by crayfish across all periods, with frequencies of occurrence close to 1 (Fig 4b). Before harvest, insects were the second most frequent food item in their diet (FO = 0.58). During the harvest period, rice took this position (FO = 0.50), and during ploughing, mammals became the second most frequent food item (FO = 0.30). The number of crayfish detected per pellet did not change with the onset of harvesting but significantly decreased during ploughing (Table 1, Online Resource 4d).

### 3.3. Food availability

A total of 25 cores (spanning 5 different paddies) and 240 crayfish transects (spanning 48 different paddies) were completed across six sampling occasions before, during and after harvesting events (day -1, 0, 1, 2, 7 and 14) (Fig. 5a). Crayfish availability in the fields increased significantly (Online Resource 3e) immediately after harvesting but then declined sharply, with almost no crayfish detected after one week. Rice grain density also increased significantly after harvest (Online Resource 3d), followed by a decline. However, unlike crayfish, rice decreased more gradually over time and remained relatively abundant even two weeks after harvest, still exceeding before harvest levels. The abundance of other food items did not change significantly due to harvesting events (Fig. 5a, Online Resource 3 f-h).

**Fig 5.**
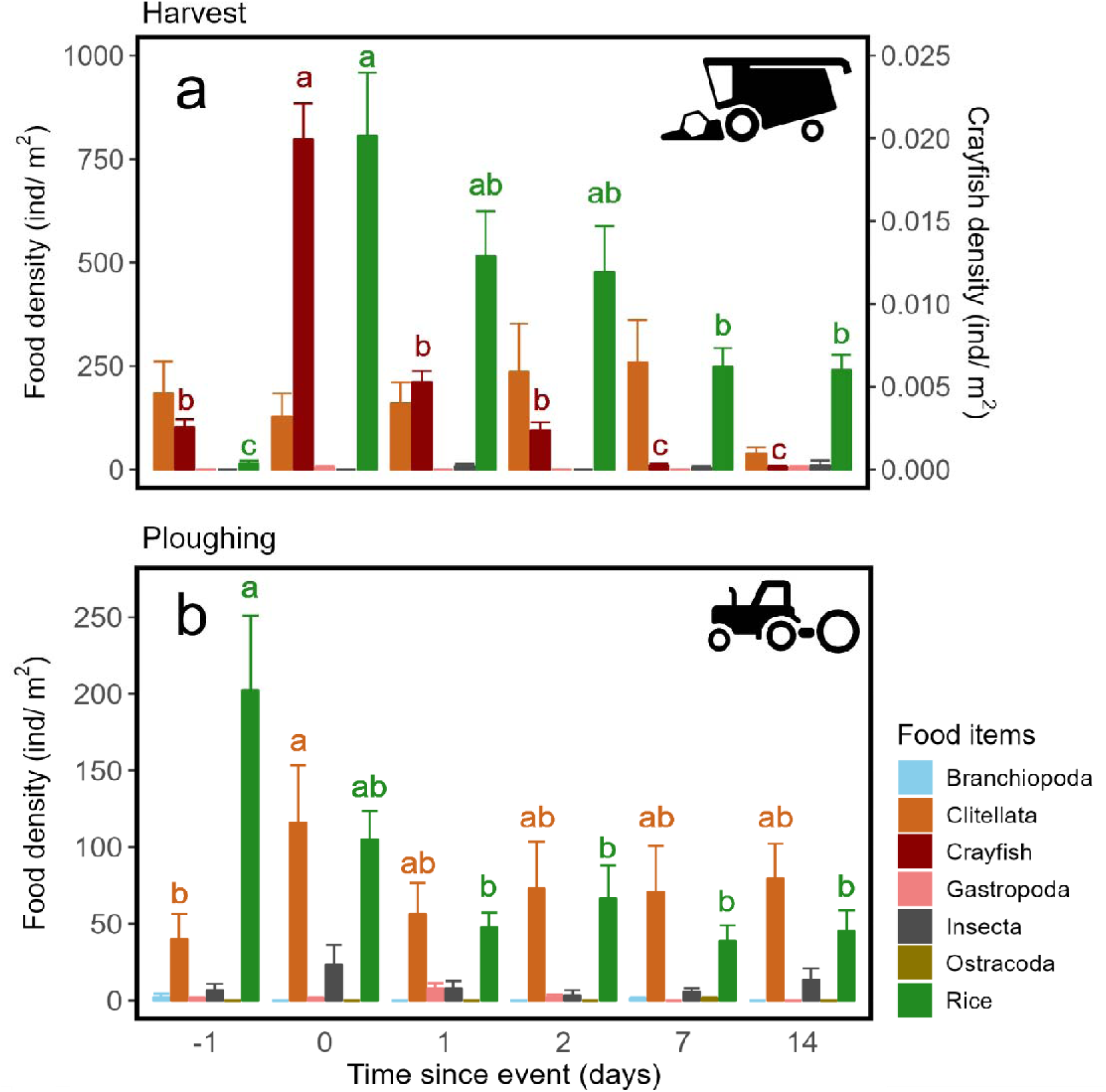
Food availability. Mean density of potential food items in flooded rice paddies before, during, and after management events. Error bars represent the standard error (SE). a – harvesting (n=25 cores; 240 transects); b – ploughing (n=80 cores; 80 transects) in flooded fields. On the y-axis, “day 0” refers to the count performed immediately after the management event (harvesting or ploughing) has finished. Differences in the mean density of food items among sampling days were assessed using GLMs with negative binomial family followed by Tuckey post hoc tests (with adjusted *P-*values for multiple comparisons) and significant differences (*P*<0.05) are depicted by different letters. Only the letter labels for food items with significant differences are shown. Note the two different y-axes in 5a depicting density of food sources sampled trough cores (left) or through walking transects (crayfish, right).

During ploughing, a total of 80 core samples and 80 crayfish transects were conducted for each of the six sampling days, encompassing 16 different rice paddies (Figure 5b). Crayfish was not detected through transects or core sampling. The two most abundant food items available were rice and worms (Class: Clitellata), while other food items were present in very low densities. Rice availability did not change with the ploughing event, but decreased significantly in the first day after ploughing, then stabilizing at relatively constant levels for the next two weeks (Figure 5b, Online Resource 3i). The abundance of worms in the soil increased significantly on the day of the ploughing event compared to the previous day (Online Resource 3j). In all subsequent periods, it remained stable, showing no significant difference from either day -1 or day 0 (Figure 5b). The abundance of other food items was not significantly affected by ploughing events (Figure 5b, Online Resource 3 k-l).

Penetrability was significantly higher in flooded fields than in dry fields, regardless of management type (Fig. 6, Online Resource 6). Among flooded fields, penetrability was highest in ploughed fields and lowest in harvested fields. However, within harvested fields, penetrability was significantly lower in ruts compared to non-rut areas. Notably, penetrability measurements in non-rut areas of flooded harvested fields were similar to those in flooded fields with rice growing. The presence of ruts in harvested fields reduced the overall penetrability of those fields (Fig. 6).

**Fig 6.**
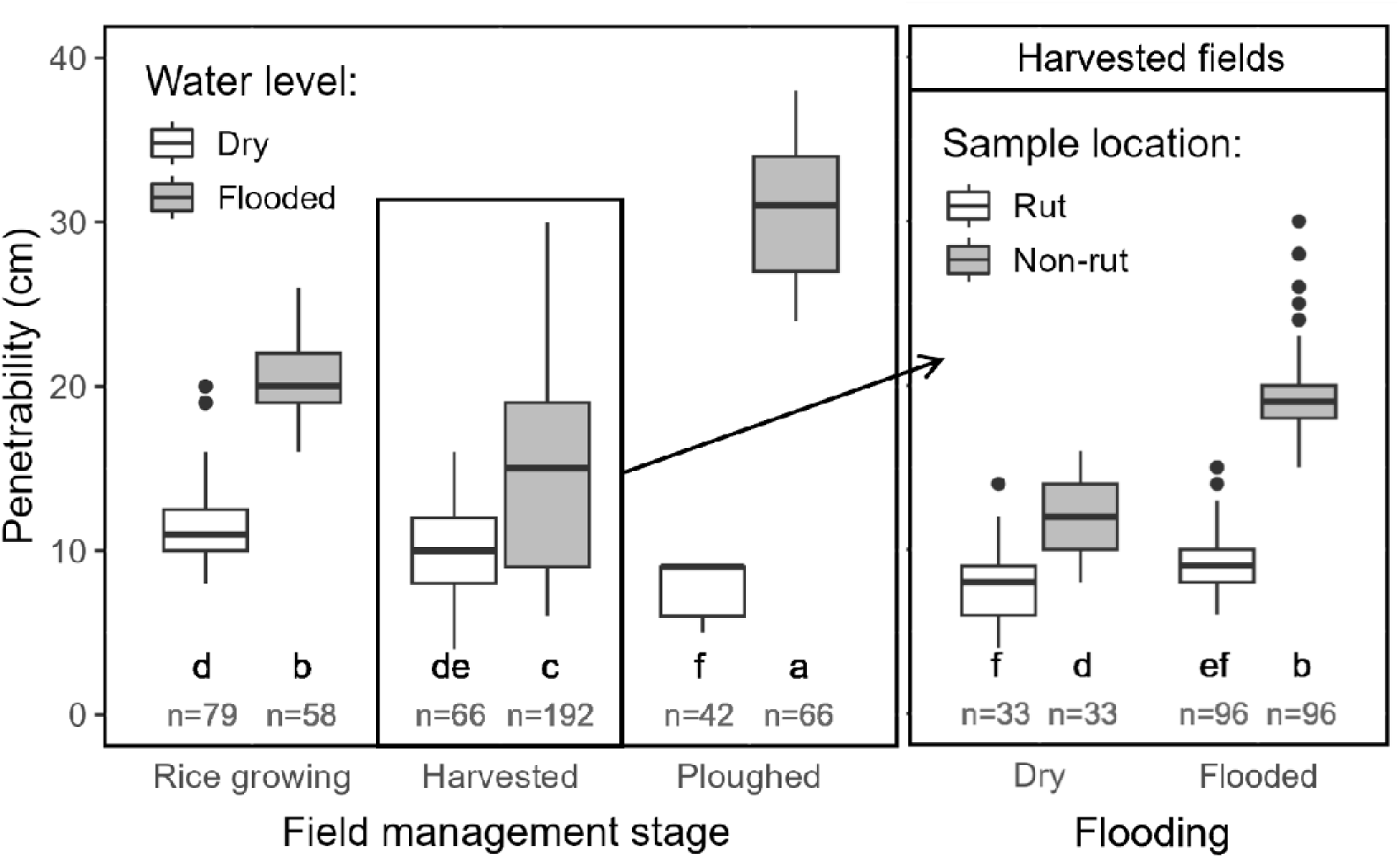
Soil penetrability of rice fields during different management stages (rice growing, harvested or ploughed) and flooding regime (flooded or dry). Soil penetrability measured in rut and non-rut locations of flooded and dry harvested fields is shown in detail. Differences in the mean penetrability among field managements were assessed using GLMs with negative binomial family followed by Tuckey post hoc tests (with adjusted *P-*values for multiple comparisons) and significant differences (*P*<0.05) are depicted by each box plot not sharing the same letter.

### 3.4. Foraging behaviour and performance, activity budgets and composition of mixed-species flocks

The foraging performance and vigilance behaviour of glossy ibis and white stork showed different patterns along the five sampling periods (Fig. 7). For both the glossy ibis and white stork, neither intake rate (prey per minute) nor success rate (proportion of successful attempts) were significantly different in birds foraging in fields undergoing harvesting or ploughing events and birds foraging during the subsequent periods in fields already harvested or ploughed (Fig. 7a,b, c and d, Online Resource 4g, j, Online Resource 5a, c). The foraging effort (steps per minute) of the glossy ibis varied across all periods during and after harvest events, but without a clear pattern (Fig. 7e, Online Resource 4h). In contrast, the foraging effort of the white stork was significantly lower in birds foraging during active harvest events (Fig. 7f, Online Resource 4j). Concerning the time spent on vigilance while foraging, neither glossy ibis nor white stork exhibited any difference between management events and the respective following stages (Fig. 7g and h, Online Resource 5b, d)”.

**Fig 7.**
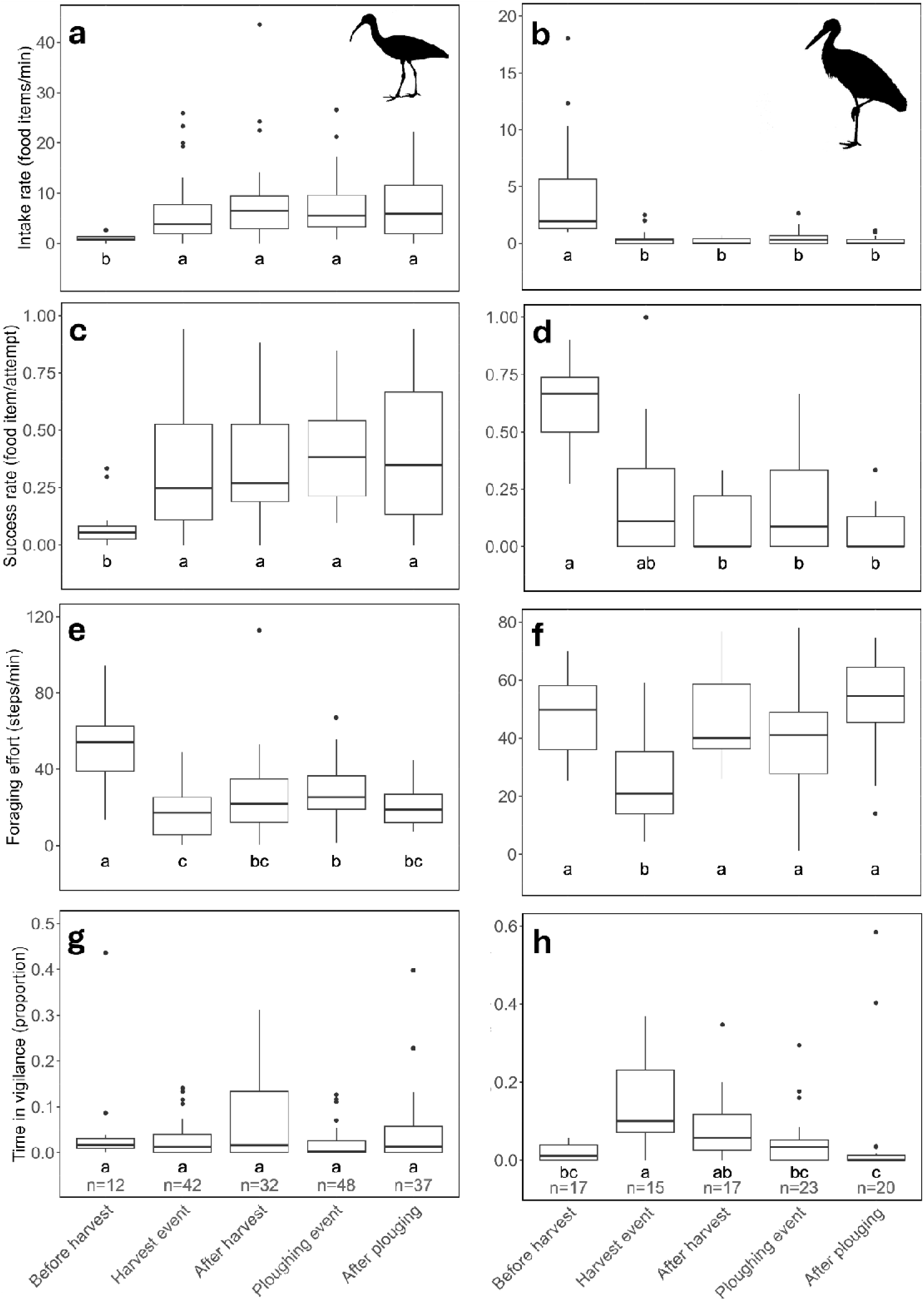
Foraging parameters of glossy ibis (a, c, e, g) and white stork (b, d, f, h) using rice fields before, during and after managing events. “Before harvest” stage corresponds to late rice-growing season, during which no fields were harvested (July to late September). The “harvest event” and “ploughing event” refer to the period up to two days following the events, while the “after harvest” and “after ploughing” stages represent periods beginning at least three days after the events. The significance of the difference in the mean of each variable across different management stages was assessed using GLMs (with a negative binomial family for non-proportion parameters and a binomial family for proportion parameters), followed by Tukey post hoc tests (with adjusted *P-*values for multiple comparisons). Groups not sharing letters are significantly different from each other (*P*<0.05).

Both glossy ibises and white storks spend significantly more time foraging in fields undergoing harvesting and ploughing compared to subsequent periods, i.e., fields already harvested or ploughed (Fig. 8a and b, Online Resource 5e, f). The size of mixed-species flocks containing either glossy ibis or white stork did not differ among management stages (Fig. 8c, d, Online Resource 4k, n), nor did the number of glossy ibis in mixed-species flocks (Fig. 8e, Online Resource 4l). It is important to note, however, that flocks were less frequently encountered in rice fields after harvest or ploughing, resulting in smaller sample sizes for these management stages. White stork’s numbers in mixed-species flocks were significantly higher during management events (both harvest and ploughing) in comparison to the following periods (Fig. 8f, Online Resource 4o), and the same patterns was recorded for the number of heterospecifics within mixed-species flocks containing glossy ibis (Fig. 8g, Online Resource 4m). For mixed-species flocks containing white stork, the number of heterospecifics showed no differences among management stages (Fig. 8h, Online Resource 4p). For details on the identity and abundance of heterospecifics across the different management stages see Online Resource 8.

**Fig 8.**
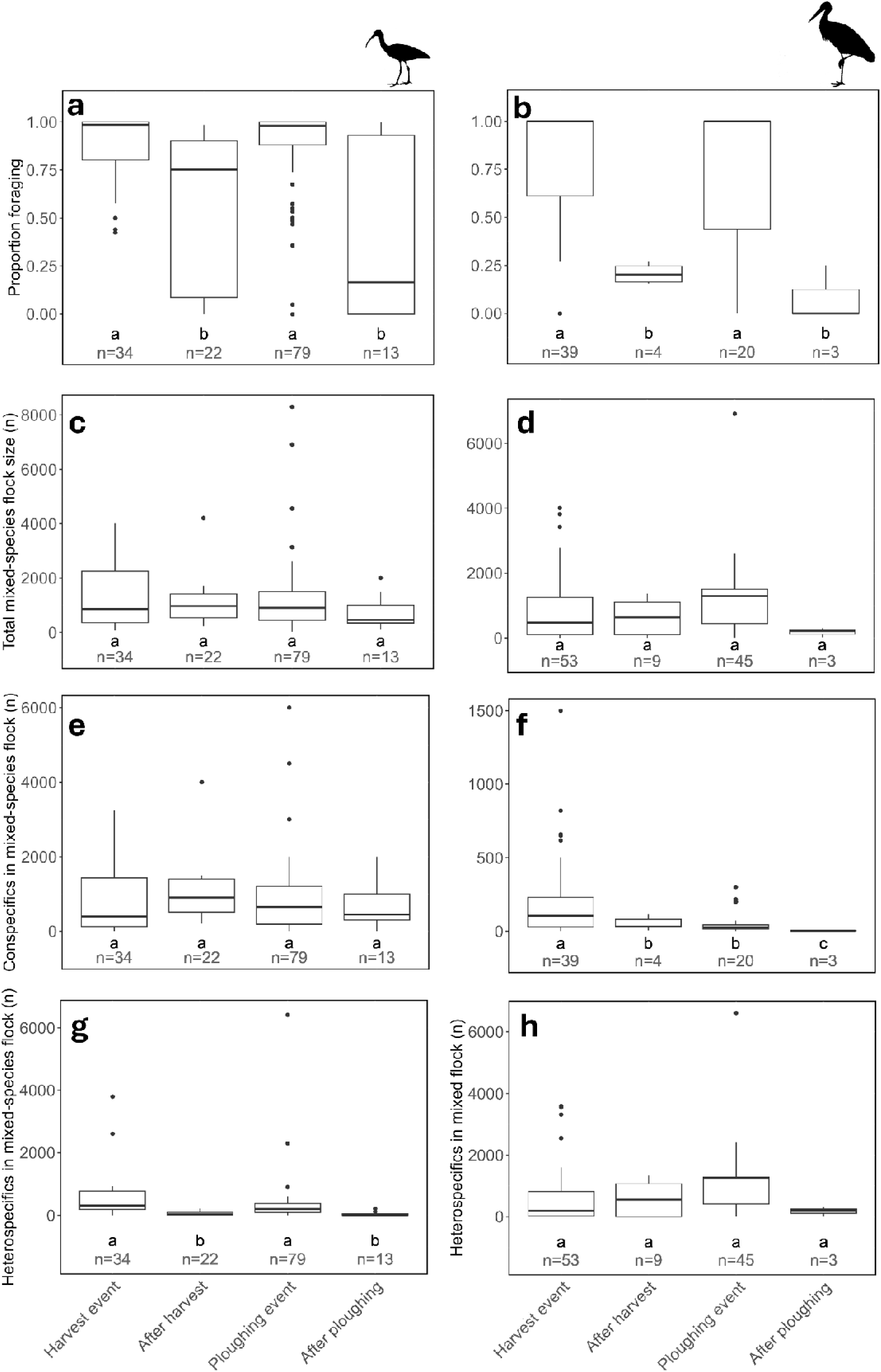
Activity budgets and flocking parameters. Activity budgets, expressed as proportion of individuals foraging, of glossy ibis and white stork and composition parameters of mixed-species flocks containing either glossy ibis (a, c, e, g) or white stork (b, d, f, h) in rice fields before, during and after managing events (harvest and ploughing). The harvest and ploughing events refer to the period up to two days following the respective events, while the after harvest and after ploughing stages represent periods beginning at least three days after the events. The significance of the difference in the mean of each variable across different management stages was assessed using GLMs (with a negative binomial family for non-proportion parameters and a binomial family for proportion parameters), followed by Tukey post hoc tests (with adjusted *P-*values for multiple comparisons). Groups not sharing letters are significantly different from each other (*P*<0.05).

## 4. Discussion

### 4.1. Waterbirds show associations with harvesting and ploughing events in rice fields

Waterbirds were shown to be strongly attracted to rice fields under harvesting and ploughing activities, creating multi-specific yet short-lived aggregations. Bird abundance decreased by more than 80% just one day after harvesting events, and nearly 50% one day after ploughing (70% decrease by the second day). GPS tracking data of glossy ibis clearly confirm that birds select fields being harvested and ploughed, as the presence of tracked birds in rice fields with working harvesting machines and tractors was overall three times higher than expected by chance alone. Similar interactions between birds and agricultural practices have been observed in other species. For example, lesser kestrels (*Falco naumanni*) in Portuguese wheat fields during harvest (Catry et al. 2014) and white storks in Polish meadows during mowing or hay clearing (Golawski and Kasprzykowski 2021) both benefit from associations with these farming activities. In the case of kestrels, foraging time was significantly reduced, and prey intake increased due to machinery flushing out prey, particularly orthopterans. However, harvesting resulted in high prey mortality, causing kestrels to avoid stubble fields once the harvesters have left (Catry et al. 2014). Similarly, storks experienced benefits when foraging in meadows being mown as the proportion of vertebrates – an energy-rich food source disturbed by machinery – in their diet increased (Golawski and Kasprzykowski 2021). The factors driving the strong association between waterbirds and managing events in rice fields will be discussed by examining the four hypotheses proposed in the introduction: (1) increased food availability, (2) enhanced foraging success and (3) reduced time allocated to vigilance.

### 4.2. Harvesting and ploughing events strongly influence waterbird food availability

Both harvesting and ploughing events increased the abundance and/or accessibility of waterbird food items, confirming the first hypothesis of this study. Waterbirds such as gulls and storks primarily feed on crayfish during the study period, whereas ibises depend mainly on crayfish before harvest and shift to rice and crayfish during and after the event. An increase in the availability of either resource should, in theory, be attractive for these birds. During harvesting, the stems of mature rice plants are cut close to the ground (Blengini and Busto 2009). This process removes the vegetation cover in the rice paddies that aquatic animals, such as crayfish, rely on for shelter and protection when foraging underwater or on the surface (Nam et al. 2015). Consequently, the accessibility of crayfish for waterbirds increases. After harvest, many individuals are likely predated by waterbirds, while others may move to non-harvested fields or irrigation channels in seek of refuge (Beingesser and Copp 1985; Correia and Ferreira 1995). Thus, the surge in crayfish availability is extremely short-lived, as indicated by the results (≈70% decrease in availability after one day). This ephemeral availability in one of the preferred food items of most waterbirds in the study area is likely an important factor attracting birds to harvest events. Harvest events are also responsible for an increase in the availability of spilled grains of rice in the soil, as harvesters are not 100% efficient in collecting rice (Alizadeh and Allameh 2013). Rice abundance also decreased over time, in part due to depletion by foraging birds (Lourenço et al. 2010), but rice grains remained abundant along the study period.

Although rice is present in both flooded and dry fields, birds are not attracted to dry harvested fields to the same extent. This suggests that increased rice availability is unlikely to be the primary factor driving this pattern, and crayfish may play a more significant role (as they occur almost exclusively in flooded fields). It is also possible that birds experience greater difficulty handling or ingesting grains in the absence of water (Guillemain et al. 1999), leading them to avoid dry rice fields. During ploughing, there is a decrease in rice grain abundance in the soil coupled with an increase in worm (Clitellata) abundance. Thus, it seems that ploughing buries some of the spilled rice from harvest beyond the reach of waterbirds as recorded in other studies (Pernollet et al. 2017) while bringing some invertebrates over from deeper within the sediment. Also, this process integrates stubble into the soil, enhancing its porosity and, consequently, its penetrability, as observed in this study and suggested by previous research (Lourenço et al. 2010). This may likely facilitate the access to benthic food items, both rice and invertebrates, deep in the soil that could otherwise be inaccessible (Nam et al. 2015), particularly for glossy ibis. A similar situation could apply to crayfish availability, as waterbirds continue to feed on them despite their absence in transect records. It is possible that a small number of immatures, cryptically coloured individuals undetectable during transects, are displaced from their burrows by tractors (Beingesser and Copp 1985; Correia and Ferreira 1995), further increasing the appeal of ploughed fields to waterbirds.

### 4.3. Waterbird foraging performance does not improve during harvesting and ploughing events

The increase in abundance of waterbirds in fields undergoing harvest and ploughing may be a response to the prompt increase in availability and/or accessibility of their preferred food as proposed by previous studies in agricultural landscapes (Catry et al. 2014; Golawski and Kasprzykowski 2021). Yet, despite the increase in food availability during active harvest and ploughing events, the large-scale association of glossy ibises and white storks with these events did not result in higher average foraging success compared with the periods following them, i.e., in rice fields harvested and ploughed more than 2 days before, when food availability or accessibility decreases. This result does not support the second hypothesis. Inter- and intraspecific interactions within mixed-species flocks may be hindering the ability of storks and ibises to fully capitalize on the surge in food availability. During harvest and ploughing events, a significantly higher number of ibises and storks actively engage in foraging in comparison with subsequent periods within the harvest and ploughing seasons. The higher number of foraging birds may indicate that competition – through resource depletion, interference or kleptoparasitism (by gulls; pers. obs.) – is likely more intense during these management events (Winterbottom 1943; Cody 1971; Kushlan 1981; Beauchamp 2005). This factor may counteract the effect of higher food availability during management events compared with the periods after, resulting in consistent foraging performance throughout the non-growing season. Similar effects have been documented in other waterbirds competing for a limited variety of resources. For instance, larger numbers of geese have been shown to negatively impact the foraging efficiency of hooded cranes in rice fields, where both species consume rice and rice ears (Zhu et al. 2020).

The fact that foraging performance of waterbirds does not increase during management events despite the increase in prey availability may also be explained by consumers functional responses, i.e, the rate of food consumption according to food densities (Solomon 1949; Holling 1959). From harvest onward, food availability may remain so high that foraging performance reaches an asymptotic maximum as the consumer is limited by its capacity to process food and no longer able to increase intake rates despite further rises in food availability during management events. This type II functional response may be present in glossy ibises, as although rice abundance declines after management events, grains remain available in large quantities throughout the entire non-growing season. Consequently, ibises are unlikely to be significantly constrained by food availability.

### 4.4. Flocking birds associated with harvesting and ploughing events do not decrease the time allocated to vigilance

The time spent on vigilance behaviour does not decrease during harvest or ploughing, thereby rejecting our third hypothesis. Typically, in flocking birds, vigilance time decreases as the conspecific number in the flock increases (Beauchamp 1998, 2001; Beauchamp and Ruxton 2008). This phenomenon has been demonstrated in some ibis species, such as the Crested ibis, *Nipponia nippon*, where an increase in the number of conspecifics within a mixed-species flock led to reduced vigilance, whereas an increase of heterospecifics did not (Ye et al. 2017). Similarly, white storks have been shown to allocate less time to vigilance as conspecific number in the flock increases (Carrascal et al. 1990; Alonso et al. 1994). The number of ibises in a flock remained unchanged during and after harvest, and, as expected, there was also no variation in the time spent on vigilance behaviour. However, the results for white storks did not align with previous literature, as their flock size decreased from management events to the subsequent periods. Despite this decline, the time allocated to vigilance did not change significantly. These unusual results may arise due to disturbance. Bird behaviour, namely vigilance, is influenced by disturbance (Wang et al. 2015), and by the perceived risk posed by the type and abundance of local predators Click or tap here to enter text.(Morrison 2011). The frequent presence of humans may strongly modify the behaviour of both the birds and their potential predators (Wang et al. 2011; Li et al. 2017). This may partially explain the atypical results regarding storks, as the harvest and ploughing seasons represents the peak of human activity in the rice fields. During this time, numerous people and heavy machinery, including harvesters, ploughing tractors and grain carrier trucks, create significant disturbances. Even if the birds get habituated to the constant disturbance (Bishop et al. 2003), these disruptions may alter the typical behavioural responses of some bird species, even those trying to take advantage of increased food availability. Also, some studies have shown that in some instances larger flocks can increase vigilance in certain species, either due to the need to monitor nearby conspecifics – possibly for social information or competition (Zhao et al. 2019) – or due to heightened alertness and faster responses to threats driven by socially transmitted fear responses (Morelli et al. 2019) or in response to higher levels of kleptoparasitism (Gonzalez 1996).

### 4.5. Waterbirds may be lured to harvesting and ploughing events by a feast perception

The rejection of hypotheses two and three suggests that, overall, waterbirds do not gain significant benefits in terms of foraging performance or reduced time in vigilance from associating with management events in rice fields. An alternative justification for these associations regards the lure of the perceived feast associated with management events and the gregarious behaviour of most waterbirds attracted to those events in the study area. Flocks of gregarious birds (e.g. storks and ibises) feed through “local enhancement”, where an individual or group seeking food in a particular area quickly identifies a good foraging location by joining a flock already feeding (Hinde 1961; Bairos-Novak et al. 2015). The noise or visual conspicuousness of the foraging flock helps with this process (Hinde 1961; Bairos-Novak et al. 2015). As long as a flock is finding food, it remains in one place and is joined by other individuals (Hinde 1961; Bairos-Novak et al. 2015). However, when the local food supply decreases, the flock moves or disbands, with its members flying off in various directions to join other flocks they can see or hear (Hinde 1961; Bairos-Novak et al. 2015). The increased food availability during harvest and ploughing events may “trap” waterbirds in this behaviour, continuously drawing them to food-rich rice fields. While some birds may benefit, others likely do not, and some may even incur costs, possibly resulting in no net gain for the community as a whole. Once birds start gathering at a harvesting or ploughing field, if they do not worsen their foraging performance, they may remain there due to behavioural inertia, even if they do not gain benefits from the aggregation. This behaviour is viewed as a form of insurance against the occasional risk of losing a good feeding spot (Ward and Zahavi 1973; Bairos-Novak et al. 2015). If that happens, the individual can quickly move to a new feeding area, rather than starting a random search across a large region (Ward and Zahavi 1973; Bairos-Novak et al. 2015).

### 4.6. Final conclusions

In conclusion, this study provides valuable insights into the relationship between birds and ephemeral food surges, particularly those generated by agricultural management events. While extensive literature documents this relationship (Toral and Figuerola 2010; Elphick et al. 2010; Johnson et al. 2011; Navedo et al. 2013; Catry et al. 2014; Golawski and Kasprzykowski 2021), the magnitude and duration of these interactions have been rarely explored, let alone their consequences in terms of benefits for the animals and the underlying drivers of their attractiveness (but see Catry et al., 2014; Golawski and Kasprzykowski, 2021). Our findings highlight the need for a more detailed examination of these interactions. While some benefits – such as increased food availability – may seem obvious, birds may not fully capitalize on them likely due to inter- and intraspecific interactions (competition and interference) and metabolic constraints (functional responses). Therefore, it is essential to determine how frequently such associations occur without clear benefits for birds and to explore what other advantages they might derive from this behaviour. For instance, even if birds do not experience increased foraging efficiency, the reduction in time spent searching for food by joining a flock already feeding in a food-abundant location – due to local enhancement – could still provide energetic benefits (Hinde 1961; Bairos-Novak et al. 2015). Additionally, although vigilance time during foraging may not decrease, the overall risk of predation might be lower due to the intense human disturbance associated with these agricultural events (Wang et al. 2011; Li et al. 2017). However, this reduced risk might not be immediately apparent, as foraging birds may prioritize resource acquisition over vigilance, maintaining a constant level of alertness regardless of perceived risk (Beauchamp 2010). These questions warrant further investigation to fully understand the complexities of bird responses to food surges driven by agricultural practices.

This study also provides important perspectives regarding the vulnerability of waterbirds in rice fields. Animals that depend on ephemeral food surges may face negative consequences if the frequency or occurrence of such surges changes (Armstrong et al. 2016). Waterbirds in rice fields demonstrate a remarkable ability to exploit these opportunities efficiently. By associating with harvest and ploughing events in the short term and rapidly shifting to the most recent one, they extend the availability of increased food resources over time (Armstrong et al. 2016). Since these events do not occur simultaneously, birds can use the resources generated by agricultural management throughout most of the non-growing season – nearly half a year. Despite efficiently exploiting food surges, our findings suggest that these birds can also successfully use alternative foraging opportunities, such as fields that were managed earlier and are no longer experiencing a surge. This flexibility may increase their resilience to potential ecological traps – for example, if changes in the timing or availability of management events reduce the predictability of these food surges (Elphick 2015).

## Acknowledgements

We thank Associação de Benificiários da Lezíria Grande de Vila Franca de Xira (ABLGVFX), namely Rui Paixão, for the permissions and critical logistic support to work on the field. Beatriz Andrês, Inês Catry, Manuel Sampaio, Catarina Taboada, Inês Marinho, Carolina Mira and Miguel Dorotea provided help in various phases of the fieldwork. We thank the two anonymous reviewers for their time and thoughtful comments, which greatly improved the quality of this manuscript.

## Declarations

### Ethics approval

Bird capture and tagging were performed in accordance with the guidelines and regulations of Institute for Nature Conservation and Forests, Portugal (permits 252/2022, 251/2023, 636/2022/CAPT, 389/2023/CAPT).

### Consent to participate

Not applicable.

### Consent for publication

Not applicable.

### Availability of data and material

The datasets generated during and/or analysed during the current study are available from the corresponding author on reasonable request.

### Code availability

The code used during the current study is available from the corresponding author on reasonable request.

### Highlighted Student Paper

*This study reveals perception, not foraging benefits, drives bird aggregations at agricultural events*.

*Findings challenge resource-driven assumptions, highlighting perception’s role in animal movement*.

### Author Contributions

JP, JPG and TC contributed to the study conception and design. Material preparation and data collection were performed by JP, TC and EC. All authors contributed to the data analysis. The first draft of the manuscript was written by JP and all authors commented on previous versions of the manuscript. All authors read and approved the final manuscript.

---

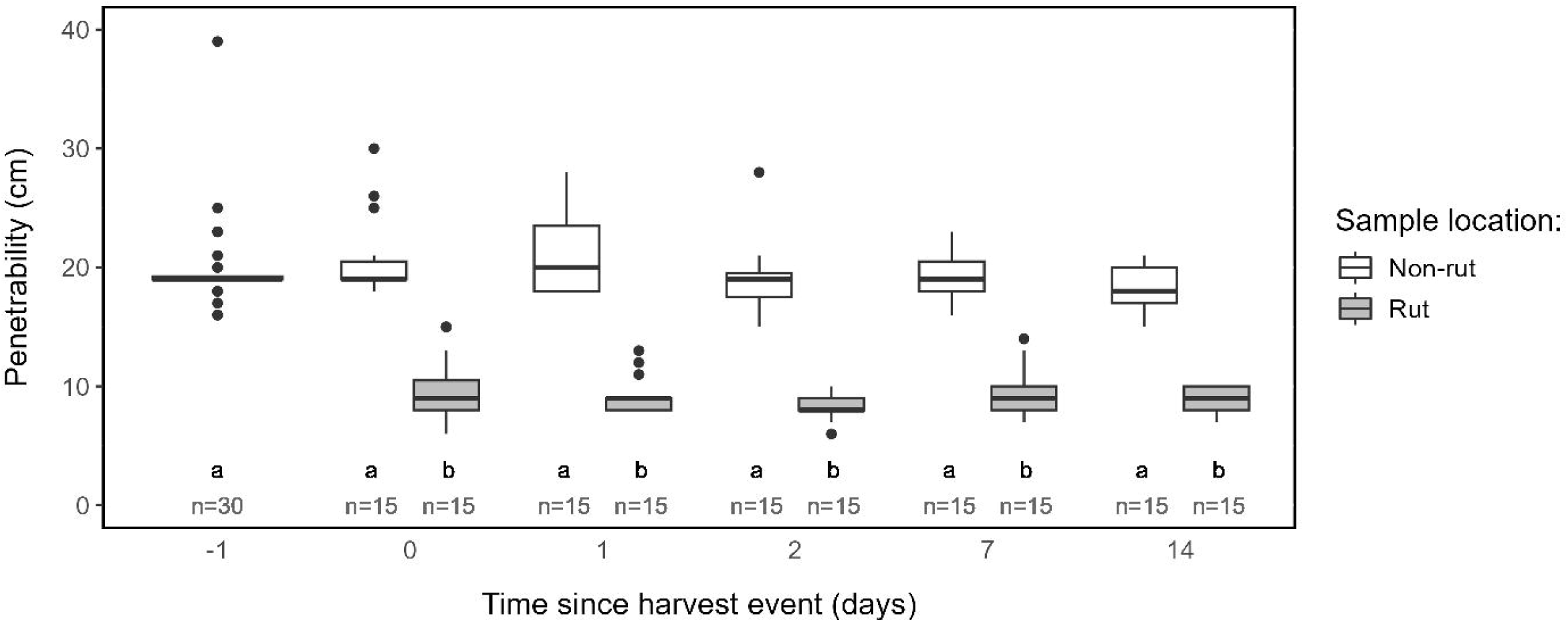

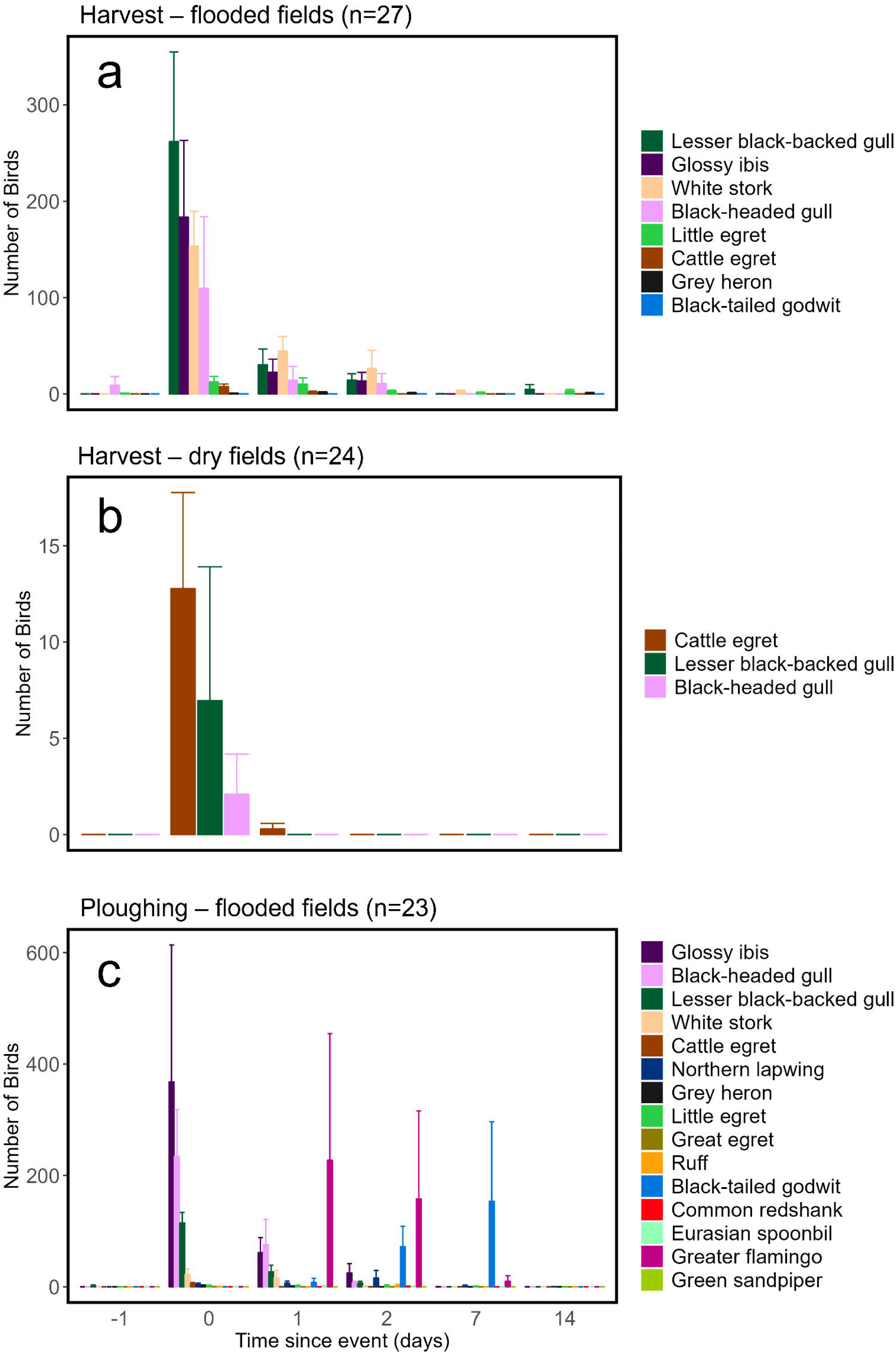

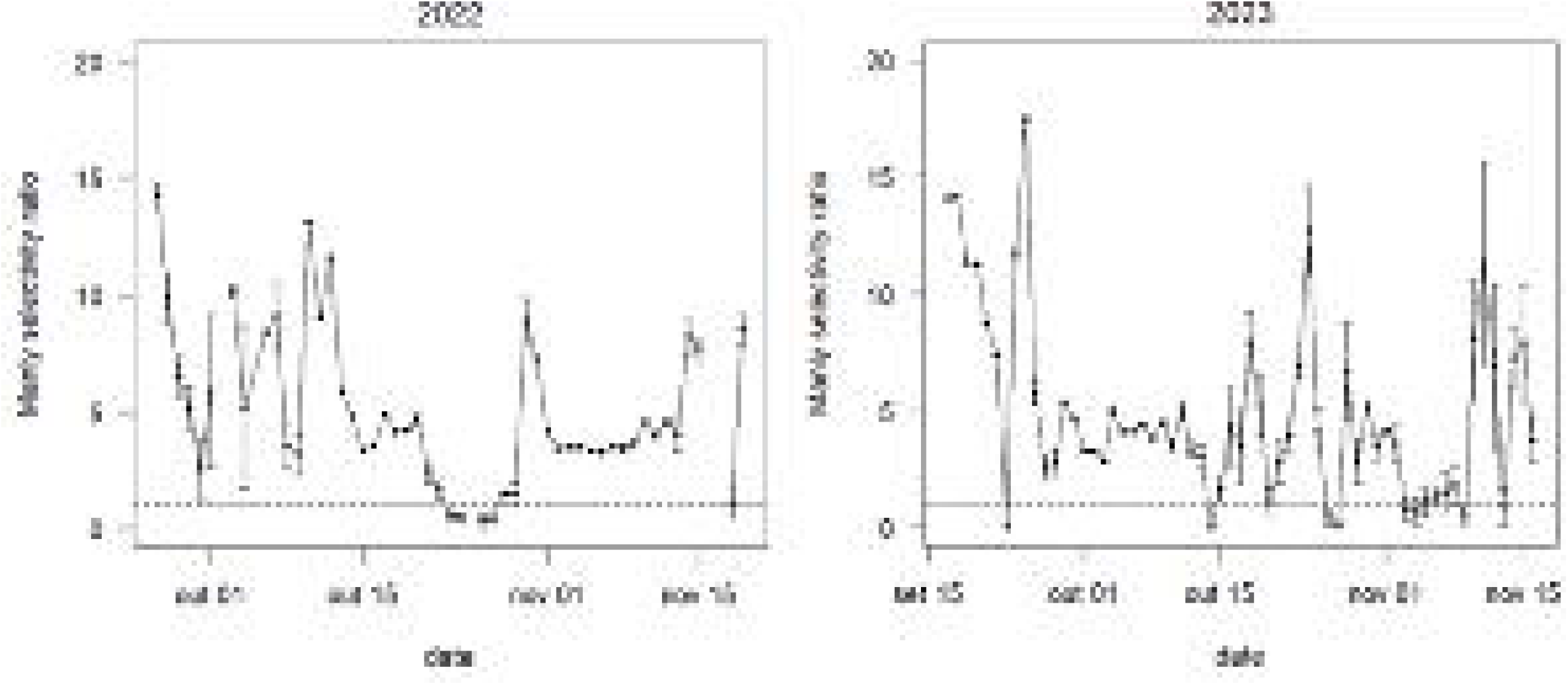

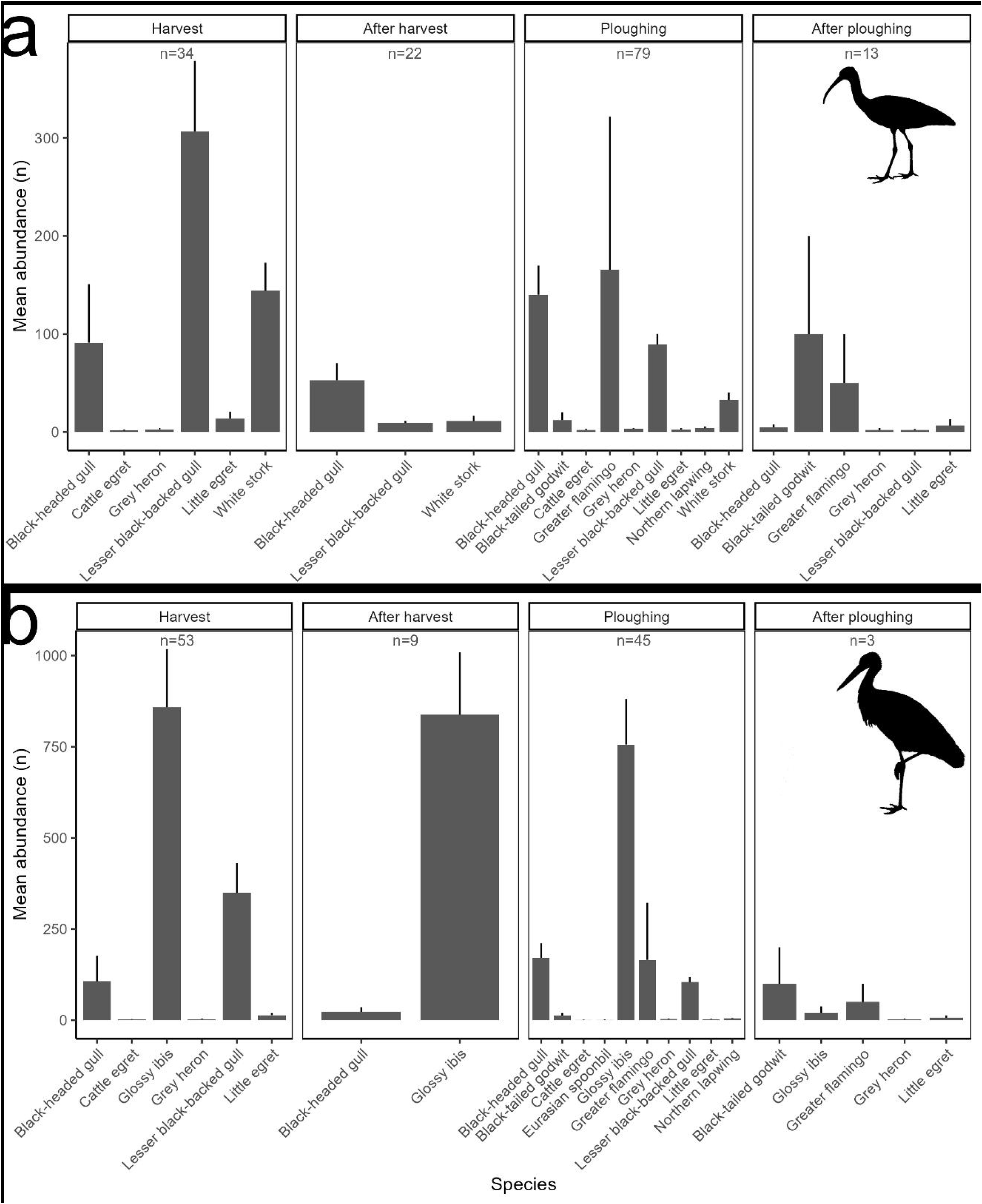

