## Supplementary Material for "Associations of waterbirds with harvesting and ploughing events in rice fields: increasing foraging opportunities or just gathering for the feast?"

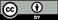
 Original content from this work may be used under the terms of the Creative Commons Attribution 4.0 license (https://creativecommons.org/licenses/by/4.0/). Any further distribution of this work must maintain attribution to the author(s) and the title of the work, journal citation and DOI.

*This research was funded in whole or in part by the Fundação para a Ciência e a Tecnologia, I.P. (FCT, https://ror.org/00snfqn58^5^) under the Grant (UID/00329/2025; LA/P/0069/2020; https://doi.org/10.54499/2020.07656.BD). For the purpose of Open Access, the author has applied a CC-BY public copyright license to any Author‘s Accepted Manuscript (AAM) version arising from this submission.*

**Supplementary Material**

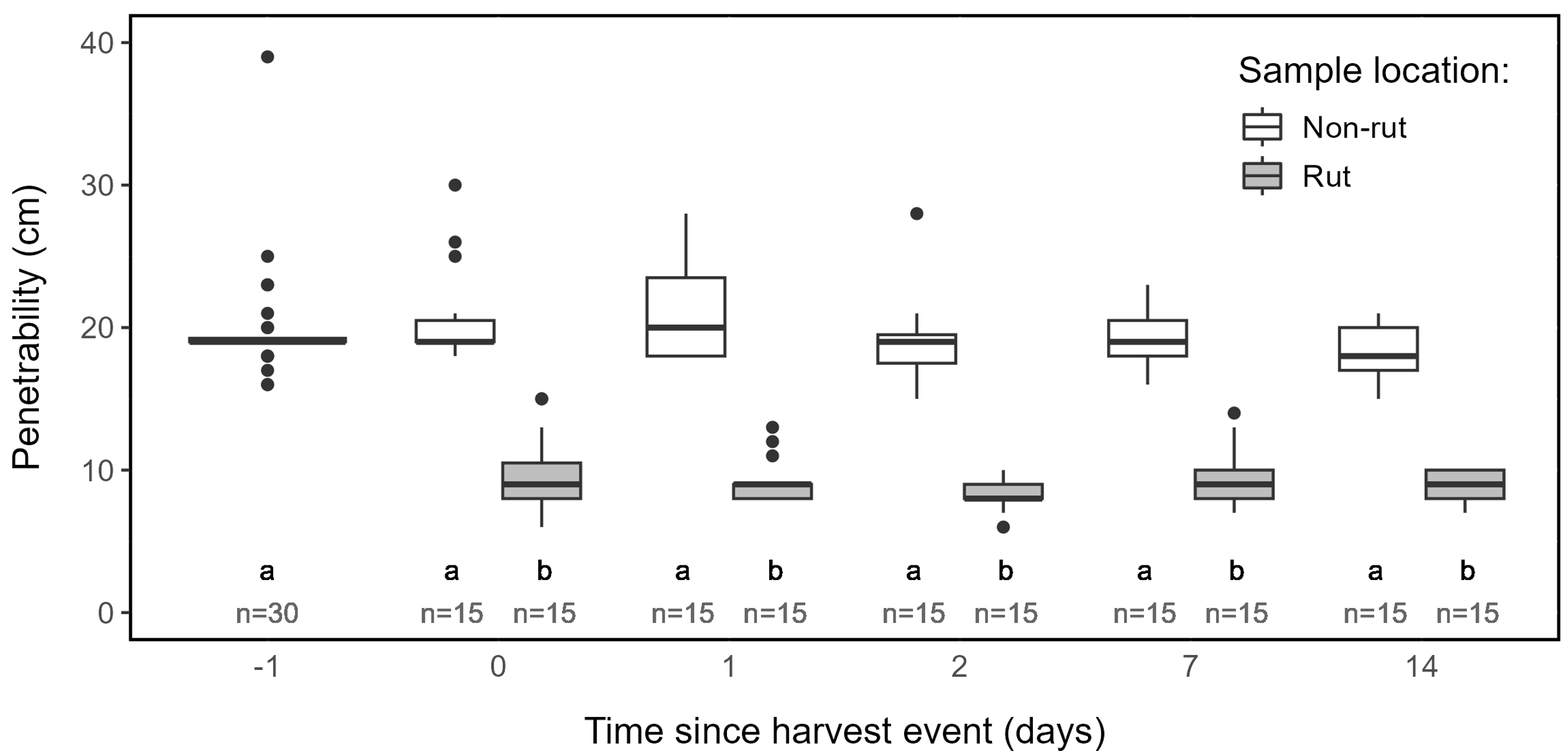

**Online Resource 1.** Soil penetrability over time for flooded rice fields subject to harvest events. On the y-axis, “day -1” refers to the day before the management event, when the field still had rice growing and “day 0” refers to the measurements taken immediately after the management action (harvesting) has finished. Differences in the mean penetrability over time were assessed using GLMs with negative binomial family followed by Tuckey post hoc tests (with adjusted *P-*values) and significant differences (*P*<0.05) are depicted by each box plot not sharing the same letter. Note the significant effect of sample location, but not time.

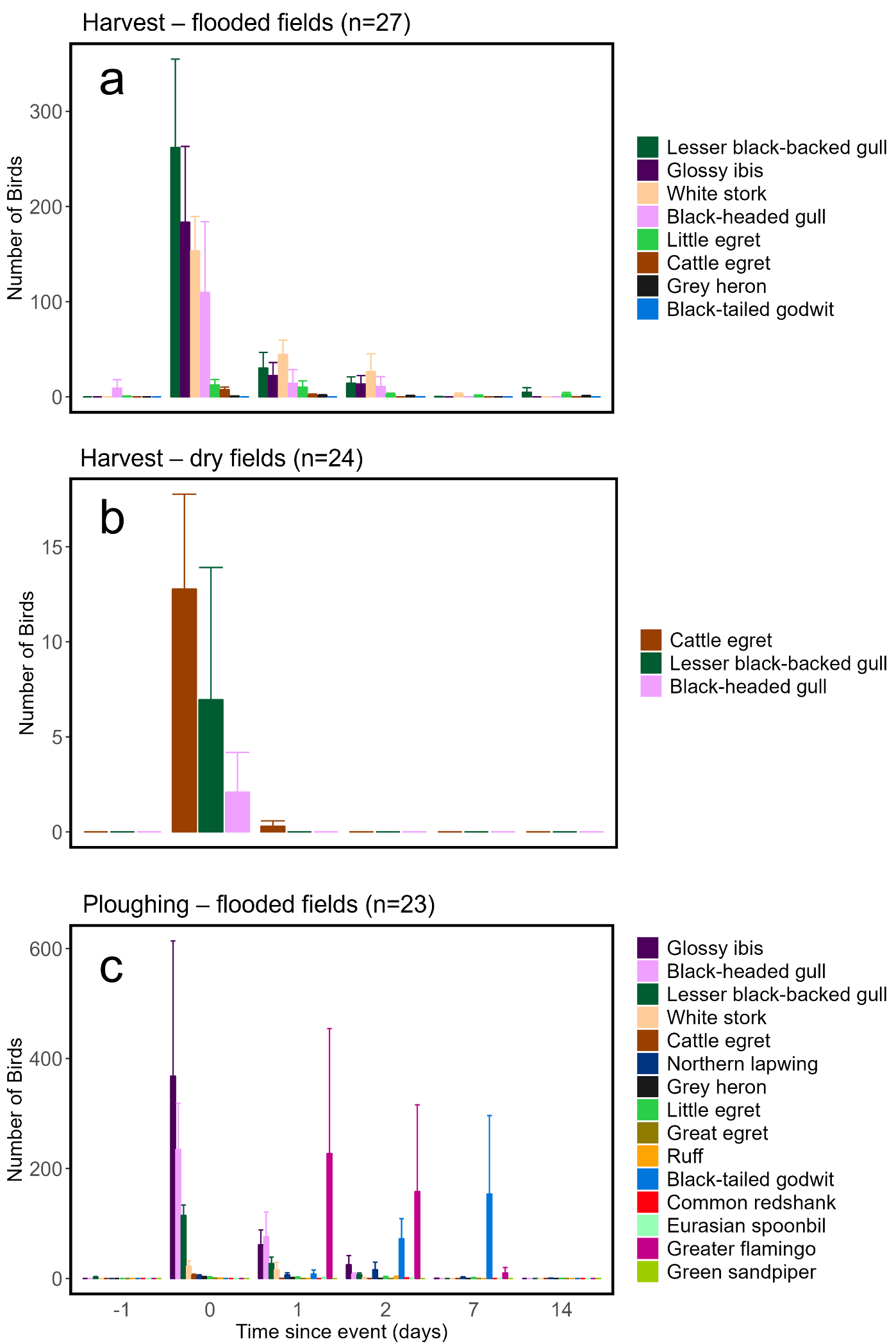

**Online Resource 2.** Mean number of waterbirds counted in rice paddies in selected days before, during, and after harvesting and ploughing events. Error bars represent the standard error (SE). a – Number of birds per species in flooded harvested fields; b – Number of birds per species in dry harvested fields; c – Number of birds per species in flooded ploughed fields. On the y-axis, “day 0” refers to the count performed immediately after the management action (harvesting or ploughing) has finished.

**Online Resource 3.** Parameters and associated statistics of generalized linear models (with negative binomial error distribution), relating waterbird abundance (a-c) or food item abundance (d-l) with the time elapsed in days since the management event (harvest or ploughing) as a factor (levels = -1, 0, 1, 2, 7 and 14 days). “Time = -1 days” was used as the reference category for all comparisons with the other factor levels. Sampling site (paddy plot identity) was used as random factor. Each model has a different response variable identified by the letters a-h. Significant effects (p-value < 0.05) are highlighted in bold.

|  | Model | | | |
| --- | --- | --- | --- | --- |
|  | Estimate | Std. error | *z-value* | *p-value* |
| **Waterbird abundance during harvest and ploughing events** |  |  |  |  |
| a) Waterbird abundance (n) - harvested flooded fields  Formula: Number of waterbirds~time elapsed to management event |  |  |  |  |
| Intercept | 2.247 | 0.535 | 4.201 | **<0.001** |
| Time=0 days | 3.778 | 0.753 | 5.000 | **<0.001** |
| Time=1 days | 2.577 | 0.779 | 3.304 | **<0.001** |
| Time=2 days | 1.985 | 0.753 | 2.633 | **0.008** |
| Time=7 days | -0.929 | 0.809 | -1.147 | 0.252 |
| Time=14 days | -0.092 | 0.757 | -0.122 | 0.903 |
| Theta, AIC, random effect variance | 0.148, 973.3, 4.505 |  |  |  |
| b) Waterbird abundance (n) - harvested dry fields  Formula: Number of waterbirds~time elapsed to management event |  |  |  |  |
| Intercept | -1.299 | 0.686 | -1.893 | **0.048** |
| Time=0 days | 4.382 | 0.881 | 4.969 | **<0.001** |
| Time=1 days | 0.067 | 0.945 | 0.071 | 0.943 |
| Time=2 days | -20.00 | 5343 | -0.004 | 0.997 |
| Time=7 days | -20.00 | 5343 | -0.004 | 0.996 |
| Time=14 days | -20.00 | 5343 | -0.004 | 0.996 |
| Theta, AIC, random effect variance | 0.107, 153.1, 0 |  |  |  |
| c) Waterbird abundance (n) - ploughed flooded fields  Formula: Number of waterbirds~time elapsed to management event |  |  |  |  |
| Intercept | 0.595 | 0.493 | 1.207 | **0.010** |
| Time=0 days | 5.394 | 0.639 | 8.433 | **<0.001** |
| Time=1 days | 4.173 | 0.633 | 6.586 | **<0.001** |
| Time=2 days | 3.901 | 0.662 | 5.887 | **<0.001** |
| Time=7 days | 2.616 | 0.648 | 4.036 | **<0.001** |
| Time=14 days | -1.000 | 0.687 | -1.455 | 0.146 |
| Theta, AIC, random effect variance | 0.300, 915.7, 0.635 |  |  |  |
| **Food availability during harvest and ploughing events** |  |  |  |  |
| d) Rice abundance (ind/m^2^) - harvest events  Formula: Density of rice grains~time elapsed to management event |  |  |  |  |
| Intercept | 2.649 | 0.323 | 8.190 | **<0.001** |
| Time=0 days | 4.043 | 0.454 | 8.897 | **<0.001** |
| Time=1 days | 3.593 | 0.456 | 7.908 | **<0.001** |
| Time=2 days | 3.519 | 0.455 | 7.743 | **<0.001** |
| Time=7 days | 2.862 | 0.455 | 6.296 | **<0.001** |
| Time=14 days | 2.833 | 0.455 | 6.233 | **<0.001** |
| Theta, AIC, random effect variance | 0.392, 1875, 0.041 |  |  |  |
| e) Crayfish abundance (ind/m^2^) - harvest events  Formula: Density of crayfish~time elapsed to management event |  |  |  |  |
| Intercept | 2.456 | 0.334 | 7.342 | **<0.001** |
| Time=0 days | 2.104 | 0.472 | 4.450 | **<0.001** |
| Time=1 days | 0.728 | 0.473 | 1.504 | 0.124 |
| Time=2 days | -0.089 | 0.473 | -0.189 | 0.850 |
| Time=7 days | -2.302 | 0.476 | -4.831 | **<0.001** |
| Time=14 days | -2.862 | 0.479 | -5.970 | **<0.001** |
| Theta, AIC, random effect variance | 0.037, 4411, 35.07 |  |  |  |
| f) Clitellata abundance (ind/m^2^) - harvest events  Formula: Density of Clitellata~time elapsed to management event |  |  |  |  |
| Intercept | -6.753 | 1.889 | -3.574 | **<0.001** |
| Time=0 days | 2.104 | 2.001 | 1.051 | 0.293 |
| Time=1 days | 0.728 | 2.301 | 0.316 | 0.751 |
| Time=2 days | -0.089 | 2.734 | -0.033 | 0.973 |
| Time=7 days | -2.302 | 6.267 | -0.367 | 0.713 |
| Time=14 days | -2.862 | 8.128 | -0.352 | 0.724 |
| Theta, AIC, random effect variance | 596.1, 53.55, 0.106 |  |  |  |

**Online Resource 3 (continued)**

|  | Model | | | |
| --- | --- | --- | --- | --- |
|  | Estimate | Std. error | *z-value* | *p-value* |
| g) Rice abundance (ind/m^2^) - ploughing events  Formula: Density of rice grains~time elapsed to management event |  |  |  |  |
| Intercept | 0.858 | 0.072 | 11.79 | **<0.001** |
| Time=0 days | 0.036 | 0.102 | 0.357 | 0.721 |
| Time=1 days | -0.466 | 0.117 | -3.979 | **<0.001** |
| Time=2 days | -0.535 | 0.119 | -4.471 | **<0.001** |
| Time=7 days | -0.661 | 0.124 | -5.302 | **<0.001** |
| Time=14 days | -0.823 | 0.131 | +6.246 | **<0.001** |
| Theta, AIC, random effect variance | 34820, 2305.6, 0.449 |  |  |  |
| h) Clitellata abundance (ind/m^2^) - ploughing events  Formula: Density of Clitellata~time elapsed to management event |  |  |  |  |
| Intercept | -0.399 | 0.136 | -2.925 | **0.003** |
| Time=0 days | 0.810 | 0.164 | 4.938 | **<0.001** |
| Time=1 days | 0.148 | 0.186 | 0.800 | 0.423 |
| Time=2 days | 0.143 | 0.186 | 0.772 | 0.440 |
| Time=7 days | 0.385 | 0.176 | 2.180 | **0.029** |
| Time=14 days | 0.562 | 0.170 | 3.442 | **<0.001** |
| Theta, AIC, random effect variance | 22704, 1924.9, 2.097 |  |  |  |

**Online Resource 4.** Parameters and associated statistics of generalized linear models (with negative binomial error distribution), relating food item abundance in droppings/pellets (a-f), foraging and vigilance parameters (g-j) or mixed-species flock parameters (j-p) with the management period as a factor (levels = before harvest, harvest, after harvest, ploughing, after ploughing; some models have only a few of these periods). “Before harvest” was used as the reference category for all comparisons with the other factor levels. Parameters that are proportions are not present in this table (See Table S3). Each model has a different response variable identified by the letters a-p. Significant effects (p-value < 0.05) are highlighted in bold.

|  | Model | | | |
| --- | --- | --- | --- | --- |
|  | Estimate | Std. error | *z-value* | *p-value* |
| **Rice and crayfish abundance in the waterbird diet** |  |  |  |  |
| a) Glossy ibis – Rice grain per dropping  Formula: Number of rice grains~management period |  |  |  |  |
| Intercept | 0.745 | 0.179 | 4.163 | **<0.001** |
| Harvest | 1.476 | 0.310 | 4.748 | **<0.001** |
| Ploughing | 1.684 | 0.284 | 5.916 | **<0.001** |
| Theta, AIC | 0.450, 864.4 |  |  |  |
| b) Glossy ibis – crayfish individuals per dropping  Formula: Number of crayfish~management period |  |  |  |  |
| Intercept | -0.048 | 0.111 | -0.436 | 0.662 |
| Harvest | -0.539 | 0.250 | -2.156 | **0.031** |
| Ploughing | -0.275 | 0.204 | -1.343 | 0.179 |
| Theta, AIC | 48800, 334.8 |  |  |  |
| c) White stork – Rice grain per dropping  Formula: Number of rice grains~management period |  |  |  |  |
| Intercept | -0.182 | 0.298 | -0.611 | 0.541 |
| Harvest | 0.484 | 0.412 | 1.176 | 0.239 |
| Ploughing | -1.368 | 0.541 | -2.529 | **0.011** |
| Theta, AIC | 0.498, 241.2 |  |  |  |
| d) White stork – crayfish individuals per dropping  Formula: Number of crayfish~management period |  |  |  |  |
| Intercept | 2.890 | 0.111 | 25.88 | **<0.001** |
| Harvest | -0.119 | 0.160 | -0.744 | 0.457 |
| Ploughing | -0.807 | 0.167 | -4.809 | **<0.001** |
| Theta, AIC | 2.541, 725.0 |  |  |  |
| **Foraging parameters** |  |  |  |  |
| g) Glossy ibis – Intake rate (food item/min)  Formula: Intake rate~management period |  |  |  |  |
| Intercept | 0.021 | 0.364 | 0.059 | 0.953 |
| Harvest | 1.771 | 0.388 | 4.556 | **<0.001** |
| After harvest | 2.065 | 0.394 | 5.233 | **<0.001** |
| Ploughing | 1.900 | 0.385 | 4.932 | **<0.001** |
| After ploughing | 1.941 | 0.391 | 4.963 | **<0.001** |
| Theta, AIC | 1.629, 986.9 |  |  |  |
| h) Glossy ibis – Foraging effort (steps/min)  Formula: Foraging effort~management period |  |  |  |  |
| Intercept | 3.977 | 0.187 | 21.19 | **<0.001** |
| Harvest | -1.122 | 0.215 | -5.223 | **<0.001** |
| After harvest | -0.717 | 0.221 | -3.239 | **0.001** |
| Ploughing | -0.690 | 0.210 | -3.278 | **0.001** |
| After ploughing | -0.975 | 0.217 | -4.476 | **<0.001** |
| Theta, AIC | 2.474, 1392.1 |  |  |  |

**Online Resource 4 (continued)**

|  | Model | | | |
| --- | --- | --- | --- | --- |
|  | Estimate | Std. error | *z-value* | *p-value* |
| i) White stork – Intake rate (food item/min)  Formula: Intake rate~management period |  |  |  |  |
| Intercept | 1.517 | 0.200 | 7.571 | **<0.001** |
| Harvest | -2.235 | 0.4558 | -4.904 | **<0.001** |
| After harvest | -3.253 | 0.633 | -5.136 | **<0.001** |
| Ploughing | -2.128 | 0.374 | -5.680 | **<0.001** |
| After ploughing | -2.806 | 0.494 | -5.672 | **<0.001** |
| Theta, AIC | 2.156, 214.06 |  |  |  |
| j) White stork – Foraging effort (steps/min)  Formula: Foraging effort~management period |  |  |  |  |
| Intercept | 3.871 | 0.111 | 34.83 | **<0.001** |
| Harvest | -0.672 | 0.166 | -4.039 | **<0.001** |
| After harvest | -0.029 | 0.157 | -0.189 | 0.850 |
| Ploughing | -0.237 | 0.147 | -1.610 | 0.107 |
| After ploughing | 0.104 | 0.150 | 0.692 | 0.489 |
| Theta, AIC | 5.286, 808.4 |  |  |  |
| **Mixed-species flock parameters** |  |  |  |  |
| k) Glossy ibis – Total flock size (n)  Formula: Number of waterbirds in flock~management period |  |  |  |  |
| Intercept | 7.182 | 0.155 | 46.27 | **<0.001** |
| After harvest | 0.022 | 0.247 | 0.092 | 0.926 |
| Ploughing | 0.013 | 0.185 | 0.072 | 0.942 |
| After ploughing | -0.578 | 0.295 | -1.959 | 0.051 |
| Theta, AIC | 1.221, 2416.8 |  |  |  |
| l) Glossy ibis – Conspecifics in flock (n)  Formula: Number of Glossy ibis in flock~management period |  |  |  |  |
| Intercept | 6.624 | 0.191 | 34.63 | **<0.001** |
| After harvest | 0.525 | 0.305 | 1.721 | 0.085 |
| Ploughing | 0.274 | 0.228 | 1.201 | 0.229 |
| After ploughing | -0.063 | 0.363 | -0.175 | 0.861 |
| Theta, AIC | 0.804, 2327 |  |  |  |
| m) Glossy ibis – Heterospecifics in flock (n)  Formula: Number of non-Glossy ibis in flock~management period |  |  |  |  |
| Intercept | 6.333 | 0.233 | 27.11 | **<0.001** |
| After harvest | -2.041 | 0.373 | -5.466 | **<0.001** |
| Ploughing | -0.497 | 0.279 | -1.779 | 0.075 |
| After ploughing | -2.892 | 0.446 | -6.473 | **<0.001** |
| Theta, AIC | 0.539, 1900.5 |  |  |  |
| n) White stork – Total flock size (n)  Formula: Number of waterbirds in flock~management period |  |  |  |  |
| Intercept | 6.831 | 0.154 | 44.29 | **<0.001** |
| After harvest | -0.449 | 0.404 | -1.111 | 0.266 |
| Ploughing | 0.282 | 0.227 | 1.239 | 0.215 |
| After ploughing | -1.655 | 0.667 | -2.480 | 0.053 |
| Theta, AIC | 0.793, 1736.3 |  |  |  |
| o) White stork – Conspecifics in flock (n)  Formula: Number of White stork in flock~management period |  |  |  |  |
| Intercept | 5.324 | 0.160 | 33.09 | **<0.001** |
| After harvest | -1.360 | 0.424 | -3.207 | **0.001** |
| Ploughing | -1.231 | 0.238 | -5.175 | **<0.001** |
| After ploughing | -4.225 | 0.769 | -5.489 | **<0.001** |
| Theta, AIC |  |  |  |  |
| p) White stork – Heterospecifics in flock (n)  Formula: Number of non-White stork in flock~management period |  |  |  |  |
| Intercept | 6.581 | 0.201 | 32.67 | **<0.001** |
| After harvest | -0.292 | 0.528 | -0.554 | 0.580 |
| Ploughing | 0.482 | 0.297 | 1.623 | 0.105 |
| After ploughing | -1.422 | 0.871 | -1.633 | 0.102 |
| Theta, AIC | 0.465, 1653.9 |  |  |  |

**Online Resource 5.** Parameters and associated statistics of generalized linear models (with beta error distribution) relating foraging and vigilance proportion parameters (a-d) or mixed-species flock proportion parameters (e-f) with the management period as a factor (levels = before harvest, harvest, after harvest, ploughing, after ploughing; some models have only a few of these periods). “Before harvest” was used as the reference category for all comparisons with the other factor levels. Each model has a different response variable identified by the letters a-f. Significant effects (p-value < 0.05) are highlighted in bold.

|  | Model | | | |
| --- | --- | --- | --- | --- |
|  | Estimate | Std. error | *z-value* | *p-value* |
| **Foraging and vigilance parameters (proportions)** |  |  |  |  |
| a) Glossy ibis – Success rate  Formula: Success rate~management period |  |  |  |  |
| Intercept | -1.735 | 0.327 | -5.308 | **<0.001** |
| Harvest | 1.098 | 0.356 | 3.077 | **0.002** |
| After harvest | 1.181 | 0.366 | 3.226 | **0.001** |
| Ploughing | 1.375 | 0.353 | 3.898 | **<0.001** |
| After harvest | 1.240 | 0.361 | 3.429 | **<0.001** |
| Phi | 3.312 | 0.443 | 7.471 |  |
| b) Glossy ibis – Proportion of time in vigilance  Formula: Proportion of time in vigilance~management period |  |  |  |  |
| Intercept | -2.578 | 0.293 | -8.774 | **<0.001** |
| Harvest | -0.390 | 0.345 | -1.132 | 0.257 |
| After harvest | 0.052 | 0.349 | 0.151 | 0.879 |
| Ploughing | -0.596 | 0.345 | -1.728 | 0.083 |
| After harvest | -0.191 | 0.339 | -0.563 | 0.573 |
| Phi | 11.99 | 2.543 | 4.718 | **<0.001** |
| c) White stork – Success rate  Formula: Success rate~management period |  |  |  |  |
| Intercept | 0.231 | 0.111 | 2.082 | **0.003** |
| Harvest | -0.730 | 0.254 | -2.877 | **0.004** |
| After harvest | -1.126 | 0.282 | -3.986 | **<0.001** |
| Ploughing | -0.776 | 0.233 | -3.328 | **<0.001** |
| After harvest | -1.361 | 0.302 | -4.501 | **<0.001** |
| Phi | 27.7 | 15.04 | 1.842 | **0.048** |
| d) White stork – Proportion of time in vigilance  Formula: Proportion of time in vigilance~management period |  |  |  |  |
| Intercept | -2.896 | 0.316 | -9.140 | **<0.001** |
| Harvest | 1.325 | 0.365 | 3.630 | **<0.001** |
| After harvest | 0.771 | 0.374 | 2.062 | **0.039** |
| Ploughing | 0.236 | 0.385 | 0.613 | 0.539 |
| After harvest | -0.496 | 0.481 | -1.032 | 0.302 |
| Phi | 8.371 | 2.034 | 4.116 | **<0.001** |
| **Mixed-species flock parameters (proportions)** |  |  |  |  |
| e) Glossy ibis – Foraging behaviour  Formula: Proportion foraging~management period |  |  |  |  |
| Intercept | 1.651 | 0.265 | 6.222 | **<0.001** |
| After harvest | -1.628 | 0.382 | -4.252 | **<0.001** |
| Ploughing | -0.053 | 0.303 | -0.175 | 0.861 |
| After ploughing | -1.990 | 0.480 | -4.145 | **<0.001** |
| Phi | 1.166 | 0.229 | 5.078 | **<0.001** |
| f) White stork – Foraging behaviour  Formula: Proportion foraging~management period |  |  |  |  |
| Intercept | 0.560 | 0.193 | 2.906 | **0.003** |
| After harvest | -1.246 | 0.621 | -2.006 | **0.044** |
| Ploughing | -0.271 | 0.324 | -0.838 | 0.402 |
| After ploughing | -2.018 | 0.693 | -2.912 | **0.003** |
| Phi | 1.514 | 0.222 | 6.792 | **<0.001** |

**Online Resource 6.** Parameters and associated statistics of generalized linear models (with negative binomial error distribution), relating penetrability with different combinations of field management stage and flooding (dry or flooded) as a factor. Significant effects (p-value < 0.05) are highlighted in bold.

|  | Model | | | |
| --- | --- | --- | --- | --- |
|  | Estimate | Std. error | *z-value* | *p-value* |
| Penetrability  Formula: Penetrability~management stage |  |  |  |  |
| Harvested flooded (ruts + non-ruts) | 0.366 | 0.043 | 8.458 | **<0.001** |
| Harvested flooded (non-ruts) | 0.671 | 0.045 | 14.85 | **<0.001** |
| Harvested flooded (rut) | -0.076 | 0.051 | -1.482 | 0.138 |
| Harvested dry (non-rut) | 0.193 | 0.063 | 3.055 | **0.002** |
| Harvested dry (rut) | -0.239 | 0.073 | -3.277 | **0.001** |
| Ploughed dry | -0.261 | 0.067 | -3.849 | **<0.001** |
| Ploughed flooded | 1.124 | 0.044 | 25.11 | **<0.001** |
| Rice dry | 0.151 | 0.050 | 2.980 | **0.002** |
| Rice flooded | 0.731 | 0.048 | 15.11 | **<0.001** |
| Theta, AIC | 14867, 4111.6 |  |  |  |

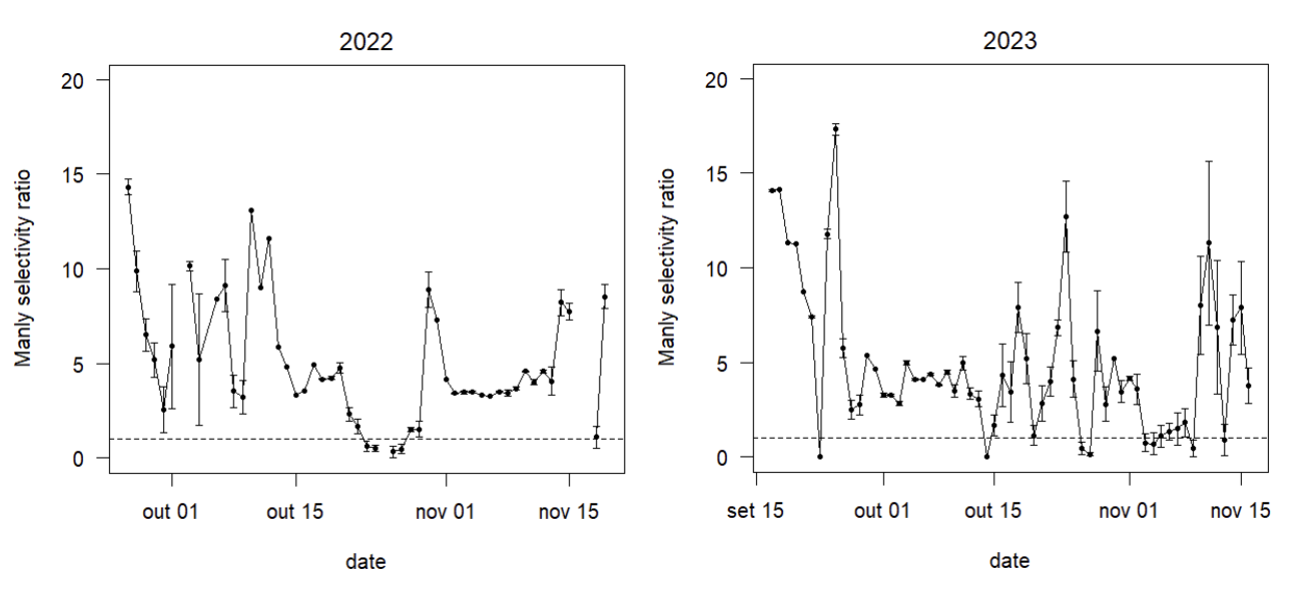

**Online Resource 7.** Manly selectivity ratios for Glossy ibis (*Plegadis falcinellus*) using rice fields undergoing management events during the harvesting season. Daily selectivity values represent the ratio between the proportion of GPS fixes in fields undergoing management events and the proportion of these fields available in each day. Selectivity ratios with 95% confidence intervals that do not overlap 1 indicate significant preference (>1) or avoidance (<1).

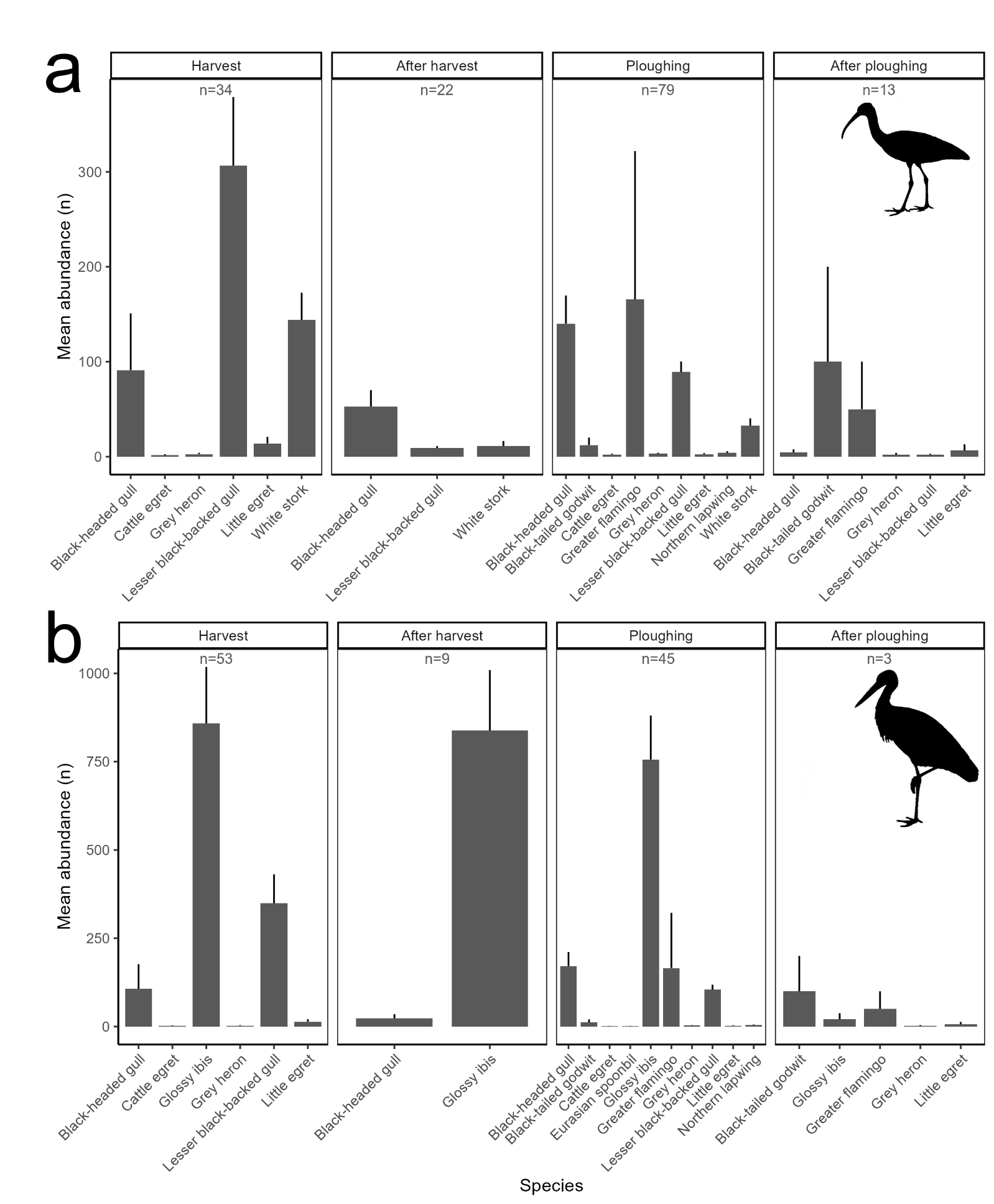

**Online Resource 8.** Mean abundance (±SE) of heterospecifics in flocks containing (a) Glossy ibis or (b) White stork in rice fields before, during, and after management events (harvest and ploughing). Only species with a mean abundance greater than 1 individual are shown. The harvest and ploughing refer to the period up to two days following the respective events, while the after harvest and after ploughing stages represent periods beginning at least three days after the events
